# Immune Profiling of Atherosclerotic Plaques Identifies Innate and Adaptive Dysregulations Associated with Ischemic Cerebrovascular Events

**DOI:** 10.1101/721688

**Authors:** Dawn M. Fernandez, Adeeb H. Rahman, Nicolas Fernandez, Aleksey Chudnovskiy, El-ad David Amir, Letizia Amadori, Nayaab S. Khan, Christine Wong, Roza Shamailova, Christopher Hill, Zichen Wang, Romain Remark, Jennifer R. Li, Christian Pina, Christopher Faries, Ahmed J. Awad, Noah Moss, Johan L.M. Bjorkegren, Seunghee Kim-Schulze, Sacha Gnjatic, Avi Ma’ayan, J. Mocco, Peter Faries, Miriam Merad, Chiara Giannarelli

## Abstract

Atherosclerosis is driven by multifaceted contributions of the immune system within the circulation and at vascular focal sites. Yet the specific immune dysregulations within the atherosclerotic lesions that lead to clinical cerebro- and cardiovascular complications (i.e. ischemic stroke and myocardial infarction) are poorly understood. Here, using single-cell mass cytometry with Cellular Indexing of Transcriptomes and Epitopes by Sequencing (CITE-seq) we found that atherosclerotic plaques were enriched in activated, differentiated, and exhausted subsets of T cells vs. blood. Next, using single-cell proteomic, transcriptomic, and cell-to-cell interaction analyses we found unique functional dysregulations of both T cells and macrophages in plaques of patients with clinically symptomatic (SYM; recent stroke of TIA) or asymptomatic (ASYM, no recent stroke) carotid artery disease. SYM plaques were enriched with a distinct CD4^+^ T cell subset, and T cells were activated, differentiated and presented subset specific exhaustion. SYM macrophages presented alternatively activated phenotypes including subsets associated with plaque vulnerability. In ASYM plaques, T cells and macrophages were activated and displayed a strong IL-1β signaling across cell types, that was absent in SYM plaques. The identification of plaque-specific innate and adaptive immune dysregulations associated with cerebrovascular events provides the basis for the design of precisely tailored cardiovascular immunotherapies.

## INTRODUCTION

Vascular inflammation is a key component of atherosclerosis that contributes to plaque instability and leads to clinical cardiovascular (CV) events including ischemic stroke and myocardial infarction^1^. Despite decades of intensive research, the immune mechanisms contributing to these processes remain largely unresolved. Most studies in the field rely on the use of animal models that fail to adequately recapitulate the human immune environment^2,3^ and lack spontaneous plaque rupture^2,4^. Pioneering histological studies from plaques of acute coronary death patients have established that culprit lesions present a large necrotic core, a thin fibrous cap and a high ratio of macrophages to vascular smooth muscle cells^5–12^. Consequently, most research has focused on macrophage infiltrates and their functionally diverse subsets as key drivers of plaque instability^13–18^. However, the inherent immune cell diversity in atherosclerotic plaques suggests a critical yet obscure role for other immune cell populations at the atherosclerotic vascular site^1,19–21^. For example, murine studies demonstrate that plaque-derived T cell subsets can be either pro- or anti-atherogenic^21–23^, but the phenotypic and functional diversity of these and other cell types in human atherosclerosis and their contribution to human plaque pathology remain undefined. Several studies support the hypothesis that circulating immune cells influence the clinical course of atherosclerosis, as patients with acute CV events present increased monocyte and CD4^+^ T cell subtypes in blood^24–31^. However, the interplay of systemic immune responses with those occurring the atherosclerotic vascular site, where rupture occurs, are vastly under investigated. Thus, immune cell phenotypes and functional relationships within and between the lesion site and the blood of the same patient are important yet unexplored concepts that could guide the design of conceptually innovative molecularly targeted immunotherapies.

Targeting inflammation to reduce CV risk in patients has long been proposed^32^, and the recent outcomes of the Canakinumab Anti-inflammatory Thrombosis Outcomes Study (CANTOS)^33^ support the clinical efficacy of this approach. However, the inhibition of the interleukin-1β (IL-1β) pathway of the innate system only moderately (−15%) reduced CV events in high risk post-MI patients, leaving a high residual CV risk. Furthermore, the failure of the Cardiovascular Inflammation Reduction Trial (CIRT)^34^ which tested low-dose methotrexate, has shown that broad anti-inflammatory treatments are ineffective in reducing CV events. Taken together these observations underscore that immunomodulatory treatments must be tailored to specific immune defects.

Therefore, the identification of the specific immune dysregulations at the plaque site could aid new viable immunotherapies for CV disease, beyond the traditional management of risk factors and use of standard of care lipid lowering drugs^35–37^. The recent use of innovative high-dimensional single-cell analyses to study murine atherosclerosis^38–40^ highlights the potential of exploiting these unbiased approaches to systematically resolve the immune diversity in human atherosclerotic plaques and decipher the molecular alterations of individual immune cells that contribute to human disease and its CV complications.

## RESULTS

### Single-Cell Immunophenotyping of Human Atherosclerosis: Study Design

To map the immune microenvironment of atherosclerotic lesions, identify mirroring immune changes in blood and pinpoint cell-specific alterations associated with clinical CV events (i.e. stoke and TIA), we performed a large-scale CyTOF mass-cytometry analysis^41^ combined with Cellular Indexing of Transcriptomes and Epitopes by Sequencing (CITE-seq)^42^ and single-cell scRNA-seq analysis of plaques from a total of 46 prospectively enrolled patients undergoing carotid endarterectomy (**Fig. 1, Supplementary Table 1a**). A first single-cell proteomic CyTOF analysis combined with an unbiased MetaClustering computational approach (cohort 1, discovery cohort; n=15) revealed an unexpected predominance of heterogeneous T cell subsets in atherosclerotic plaques compared to blood. We also identified discrete adaptive dysregulations within plaques of patients with recent cerebrovascular events (i.e. stroke, TIA) (symptomatic, SYM; n=7) vs. asymptomatic patients (ASYM; n=8) (**Supplementary Table 1b**). Based on these initial observations, we performed an in-depth CyTOF analysis of T cells in blood and plaque of the same patient (n=23) (**Fig. 1b**) in prospectively enrolled ASYM (n=14) and SYM (n=9) patients (cohort 2, validation cohort) (**Supplementary Table 1c**). We complemented these studies with Cellular Indexing of Transcriptomes and Epitopes by Sequencing (CITE-seq) analysis of blood and paired atherosclerotic plaque of the same patient. Finally, to identify plaque specific immune alterations associated with clinical cerebrovascular ischemic events, we investigated alterations of T cells and macrophages in atherosclerotic plaques of SYM and ASYM patients combining CyTOF, scRNA-seq and cell-cell interaction analyses (**Fig. 1c**).

**Figure 1.**
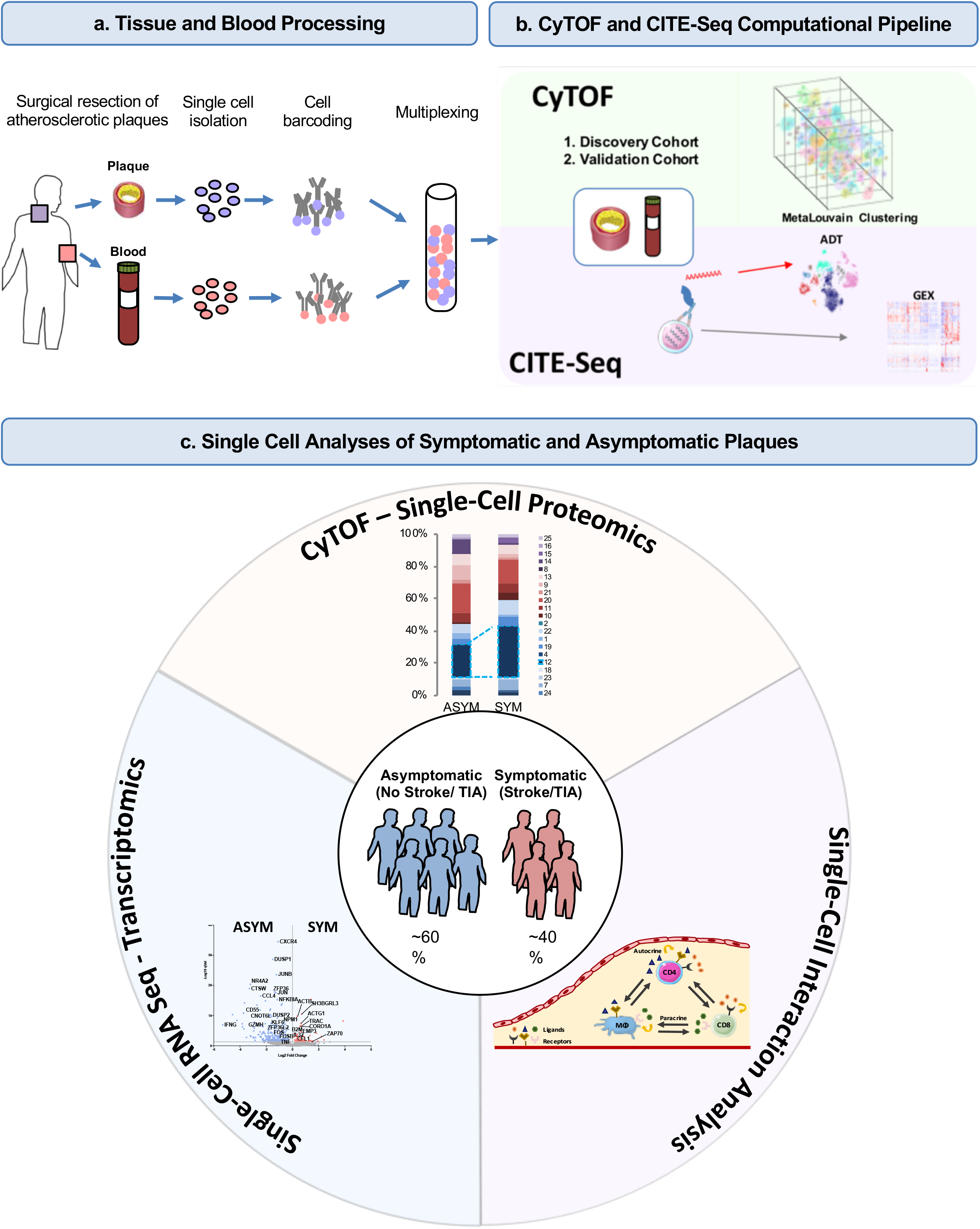
Study Overview. (a) Collection and processing of atherosclerotic tissue (plaque) and paired blood from the same patient. (b) Comparison of immune cells in atherosclerotic tissue and paired blood using CyTOF or CITE-seq analyses. (c) Single-cell analysis of immune dysregulations associated with cerebrovascular events (i.e. Stroke, TIA) in symptomatic (recent stroke) and asymptomatic (no recent stroke) patients with carotid artery disease: a three tier approach. Analysis of: 1. single-cell proteomic data from CYTOF, 2. Gene expression data from scRNA-seq, and 3. Cell-Cell communications based on ligand and receptor interactions. ADT: Antibody-tag data from CITE-seq; GEX: gene expression data from CITE-seq.

### Single-Cell Mass-Cytometry Immunophenotyping of Atherosclerotic Plaque and Paired Blood

The phenotypic distribution of immune cells in atherosclerotic tissue and blood by CyTOF were analyzed using viSNE^43^, an unbiased computational tool that distills high-dimensional single-cell data into two dimensions (**Fig. 2a**). Unbiased MetaLouvain clustering analysis^44,45^ was then used to first cluster single cells by shared proteins in each tissue and blood for individual patient. Then, secondary clustering identified MetaClusters (MCs) common across tissues and patients. MCs discovered in cohort 1 (discovery cohort, n=15) largely corresponded to known immune cell populations with distinct distributions between blood and plaque of patients (**Fig. 2a-d, Extended Fig. 1a and b**). All major immune cell lineages were identified in atherosclerotic lesions, with CD4^+^ and CD8^+^ T cells combined being the most abundant (≈65%), and with CD8^+^ T cells being the most enriched vs. blood (**Fig. 2d**). Signatures stratified by tissue type, identified immune cells specifically enriched in atherosclerotic lesions that included macrophages (MC 4), CD8^+^ EM T cells (MC 12), CD4^+^ EM T cells (MC 7) and CD4^−^CD8^−^ T cells (MC 18) (**Fig. 2e, f, Extended Fig. 1b; Supplementary Table 2**). As expected, CD14^+^ monocytes (MC 13) and natural killer (NK) cells (MC 6) were more abundant in blood. Similarly, plasmacytoid dendritic cells (pDCs) (MC 16), B cells (MC 3), and CD4^+^CD8^+^ T cells (MC 10) were enriched in blood, although the latter accounted for < 3% of the overall immune compartment (**Extended Fig. 1b and c**).

**Figure 2.**
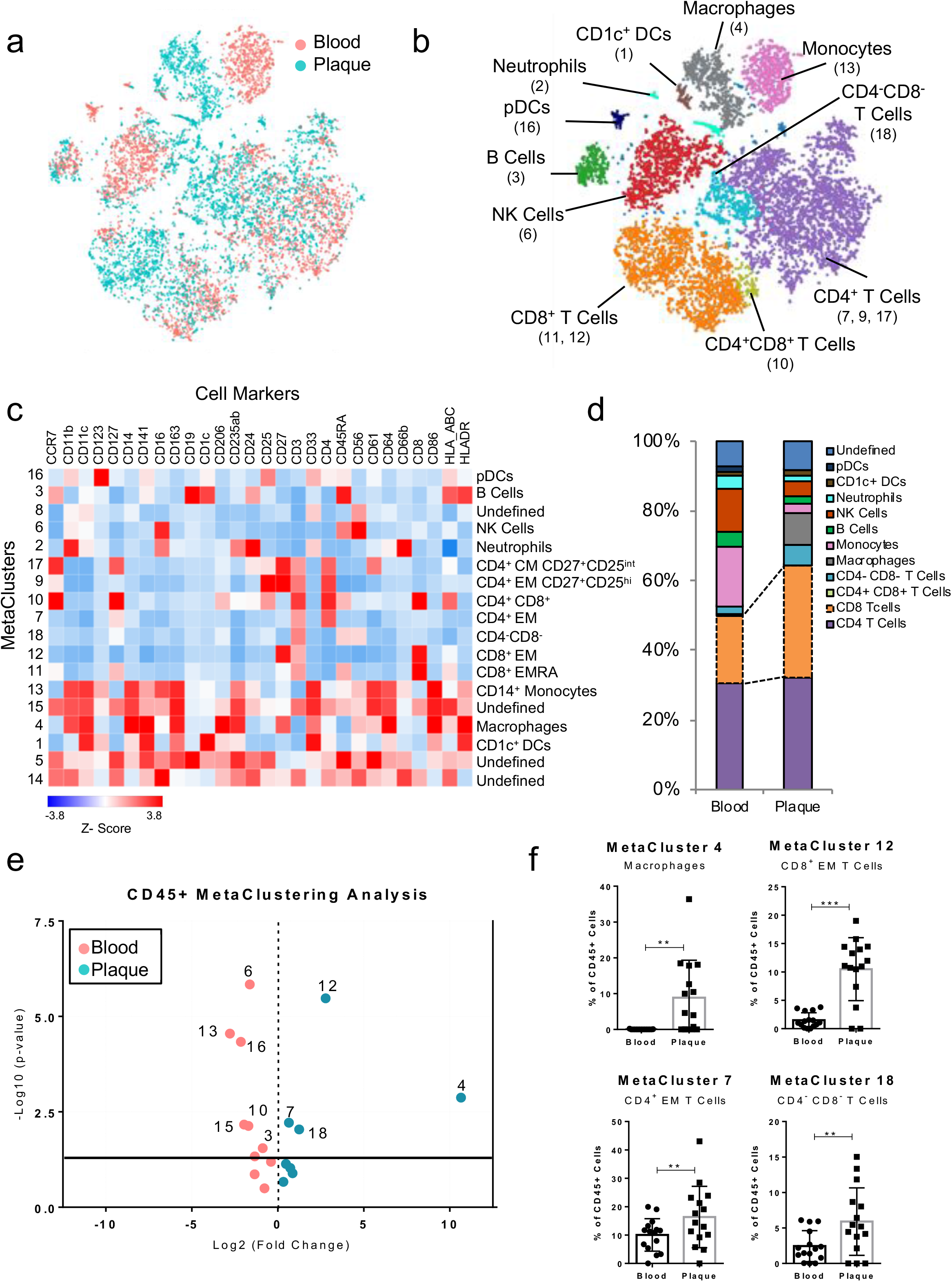
T Cells and Macrophages Dominate the Atherosclerotic Immune Landscape. (a, b) MetaLouvain Clustering of CD45^+^ cells derived from blood and atherosclerotic plaque tissue (n=15). Representative ViSNE plots of immune cells displaying tissue type (a), or overlaid with color-coded immune cell populations (b). (c) Heatmap of normalized, Z-scored protein expression of MetaCluster (MC) data, relating the MC communities (left) to the expression of protein markers (top) and the annotated cell types (right). (d) Bar chart of the relative frequency of immune cell types derived from aggregated MC data. pDC, plasmacytoid dendritic cell; DC, dendritic cell. (e) Volcano plot of the MC frequency fold change atherosclerotic plaque, color coded by the tissue enrichment (plaque: blue, blood: pink). P values were calculated using a paired, two-sided Student’s t-Test and FDR calculation in *R*. (f) Scatter bar plots of population frequency from MCs enriched in plaque tissue. Data were analyzed with the Wilcoxon test. Values are mean ± SD **p<0.01, ***p<0.001.

### Single-Cell Mass-Cytometry of Immune Cells in Atherosclerotic Plaque

An independent MC analysis comparing immune cells derived from ASYM (n=8) and SYM (n=7) plaques from patients of cohort 1 identified 15 plaque tissue-specific MCs (**Extended Fig. 2a-d**). These included 2 MCs of macrophages (MC 1 and MC 4). In particular, macrophages of MC 1 showed high expression of CD206 and CD163 (**Extended Fig. 2a**), markers of alternative M2-like macrophages in atherosclerotic tissue that have been associated with either atheroprotective or proatherogenic functions^46,47^. This analysis also revealed 9 MCs of T cells in plaques which comprised 3 subsets of CD8^+^ T cells (MCs 3, 7 and 10) and 5 of CD4^+^ T cells (MCs 5, 8, 9, 13 and 15) (**Extended Fig. 2a, c, d, see Supplementary Information and Supplementary Fig. 1**). Interestingly, we found an exclusive enrichment of CD8^+^ T cells (MC 7) and of CD4^+^CD56^+^ cells (MC 5) in SYM plaques (**Extended Fig. 2e**). Remarkably, these differences were not driven by plaque pathology—as both prospectively enrolled ASYM and SYM patients of cohort 1 presented the same type VI plaque according to the American Heart Association (AHA) classification^47^—nor by their clinical characteristics (**Supplementary Information and Supplementary Fig. 2a-e**).

### Single-Cell Mass-Cytometry Analysis of T Cells in Atherosclerotic Plaque and Paired Blood

The findings from our discovery cohort (cohort 1) suggested that in human atherosclerotic plaques T cells were the most abundant type of immune cell, displayed a greater heterogeneity than macrophages and presented unique alterations associated with clinical CV events. To validate these data, further resolve the heterogeneity of the T cell compartment in human atherosclerosis, and identify new specific adaptive dysregulations in plaques, we analyzed a second independent cohort (cohort 2, validation cohort, n=23) of prospectively enrolled ASYM and SYM patients (**Supplementary Table 1**) with an extended mass-cytometry antibody panel (**Fig. 1a**; **See Methods and Supplementary Table. 3**). This analysis confirmed that T cells were dominant in plaques compared to blood, and discovered an unprecedented diversity of the adaptive immune compartment in atherosclerosis (**Fig. 3a-d, Extended Fig. 3a-b**). We identified thirteen MCs of CD4^+^ T cells that included central memory (CM) (MCs 7, 14, 17, 23 and 24), effector memory (EM) (MCs 1, 4, 12, 18, and 22), terminally differentiated effector memory (EMRA) (MC 2), and regulatory T cells (MCs 6 and 19). The CD8^+^ T-cell compartment comprised 6 MCs, including naïve (MC 13), EM (MCs 20, 11 and 10), and EMRA (MCs 9 and 21) (**Fig. 3e, Extended Fig. 3a-c; See Supplementary Information and Supplementary Fig. 3**).

**Figure 3.**
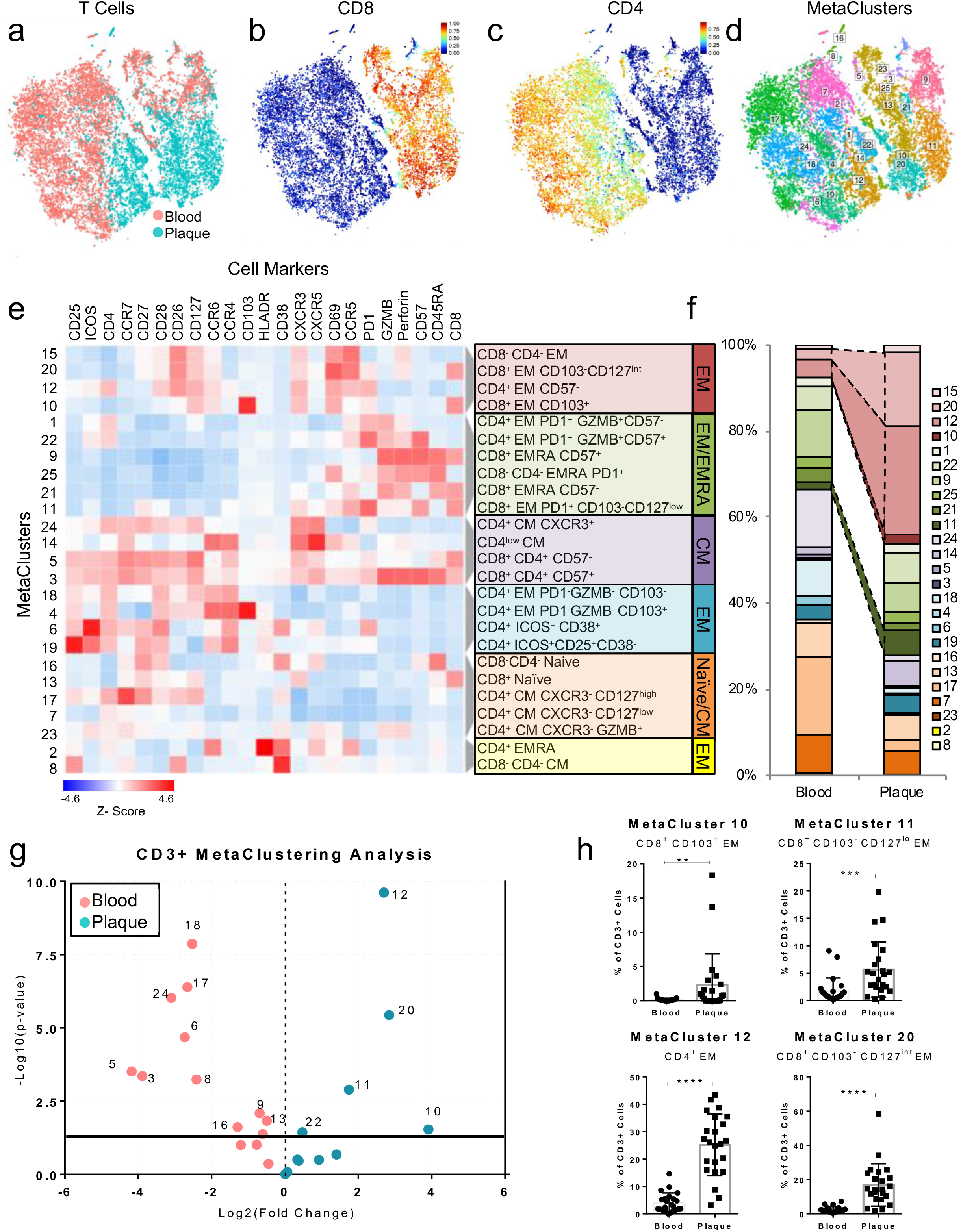
Diversity of the T-Cell Compartment in Human Atherosclerosis. (a-d) MetaLouvain clustering of T cells from blood and plaque tissues (n=23). Representative viSNE plot shows the distribution of blood and plaque (a), the expression of the T-cell markers CD8 (b) and CD4 (c), and the distribution of the T cell MC communities (d). (e) *Clustergrammer* heatmap showing protein marker expression (top) in each MC (left) and the canonical annotation of these communities (right). Light gray dendrogram bars indicate the clustering of MCs based on the cosine distance method in *Clustergrammer*. (f) Bar chart of the relative frequency of T-cell MCs. (g) Volcano plot of the fold change of MC frequency in blood (left) and atherosclerotic plaque (right). P values were calculated using a paired, two-sided Student’s t-Test and correspondent FDR calculated in R. (h) Scatter bar plots of population frequency from MCs enriched in plaque tissue. (**p<0.01, ***p<0.001, ****p<0.0001) Data were analyzed with the Wilcoxon test. Values are mean ± SD.

Atherosclerotic lesions had a higher frequency of CD8^+^ T cells (≈39% vs. ≈26% in blood), and a lower frequency of CD4^+^ T cells (≈50% vs. ≈65% in blood) (**Fig. 3f**), confirming our initial finding of a CD8^+^ T cell enrichment in plaque relative to blood. A cross-tissue analysis revealed 4 MCs enriched in atherosclerotic plaques (**Fig. 3g, Extended Fig.3b**) that were all EM T cells (CCR7^low^ and CD45RA^low^) and shared high expression of CD69, an early activation and tissue residency marker^48^ (**Fig. 3e**). Three of these MCs were CD8^+^ subsets (MCs 10, 11, 20), and one was a CD4^+^ subset (MC 12) (**Fig. 3e-h, Extended Fig. 3d**). Despite belonging to different T cell lineages, MCs 10, 12, and 20 displayed the most similar marker expression pattern and unbiasedly clustered together (**Fig. 3e**, **Extended Fig. 3e**), suggesting similar functional states in plaque. Conversely, MC 11 unbiasedly clustered with EMRA subsets, suggesting either functional similarities or a transition toward a terminally differentiating phenotype of this MC (**Fig. 3e, Extended Fig. 3d**). A key observation was that CD8^+^ T cells from MC 10 were CD103^+^ and detected exclusively in plaque (**Fig. 3e and h, Extended Fig. 3b**), and corresponded to a classical tissue-resident memory (TRM) T cell that has been reported in human lymphoid and non-lymphoid tissues^48–50^. Our study is the first to identify their presence in the atherosclerotic arterial wall in humans.

### Single-Cell Mass-Cytometry Analysis of T Cells Enriched in Atherosclerotic Plaques

Plaque-derived CD4^+^ and CD8^+^ EM T-cells of MCs 11, 12, and 20 expressed markedly higher levels of CD69 compared to blood (**Extended Fig. 4a-c**). Because CD69 can define TRM subsets regardless of CD103 expression^48^, these observations suggest that plaque-enriched MCs might represent TRM subsets. CD69 is also a marker of early activation; therefore, we assessed the activation markers CD38, CCR5, and HLA-DR^48,51^. CD38 was uniformly upregulated on cells of MCs 11 and 12, and MC 20 compared to blood. In contrast, CCR5 and HLA-DR showed different patterns of expression. CCR5 was upregulated in plaque-derived cells from MCs 11 and 12, while MC 20 had similarly high levels of CCR5 regardless of origin. HLA-DR was only upregulated in tissue cells of MC20 compared to blood (**Extended Fig. 4a-c**). These data suggest that CD4^+^ and CD8^+^ T cell subsets that are enriched in atherosclerotic plaque tissue are more activated than their blood counterparts and present a heterogeneous spectrum of activation.

Surprisingly, T cells from MCs 11, 12, and 20 expressed high levels of PD-1, a negative regulator of T cell activation and marker of T cell exhaustion^52–54^, in atherosclerotic tissue compared to blood. Additionally, T cells in tissue generally expressed lower levels of the co-stimulatory molecules CD28, CD27, and CD127 compared to blood (**Extended Fig. 4a-c**). Overall, plaque T cell subsets exhibited a chronically activated and differentiated phenotype, and because they presented high PD-1 levels, these subsets may simultaneously initiate a regulatory feedback and exhaustion reprogramming in response to plaque chronic inflammation^52–54^. Consistent with this observation, we found lower levels of perforin, but not of granzyme B, in plaques for CD8^+^ T cells of MC 11, 12 and 20, which suggests only partially reduced cytotoxic functions, typically seen in early phases of T cell exhaustion^55–57^ (**Extended Fig. 4d**). Because MC 10 was exclusive to atherosclerotic plaque, we compared it to the other CD8^+^ EM subsets (MCs 11 and 20) in tissue (**Extended Fig. 4c**). CCR5 expression was similar between MCs 10 and 20, and higher than in MC11, suggesting higher activation of both MC 10 and MC 20 than MC 11 in tissue. MC 10 also displayed lower levels of the co-stimulatory molecules CD27 and CD28 than MCs 11 and 20, suggesting that the cells of MC 10 were more differentiated than other EM CD8^+^ T cells in tissue^52^. Comparable levels of PD-1 between all CD8^+^ from MCs 10, 11, and 20 suggested a similar degree of exhaustion (**Extended Fig. 4c**). An independent MC analysis on plaque-derived T cells from the same cohort (cohort 2) identified a MC of CD69^+^CCR5^+^PD1^int^CD127^−^ CD8^+^ T cells whose frequencies correlated with TCR clonality in tissue, suggesting clonal expansion of these cells in atherosclerotic plaques (see **Supplementary Information and Supplementary Fig. 4**).

### Simultaneous Surface Marker and Gene Expression Immune Profiling Across Plaque and Blood: CITE-seq Analysis

In order to further investigate the identified functional differences and decipher key differences in immune molecular mechanisms and signaling pathways in plaque tissue and in blood, we analyzed the T cell compartments using Cellular Indexing of Transcriptional Epitope Sequencing (CITE-seq)^42^, an innovative method that integrates proteomic (surface marker expression) and transcriptional (gene expression, GEX) data into a single readout. We also focused on plaque macrophages, given their undisputed role in the pathogenesis of the disease^17,21,46,58,59^. We analyzed 5,232 total cells (1,643 from plaque and 3,589 from blood) and assigned cell types according to the canonical surface marker expression from the Antibody Derived Tag (ADT) data. Consistent with our CyTOF results T cells accounted for the majority of all immune cells in plaque (**Fig. 4a-c**) and CD8^+^ T cells were enriched in plaque (≈46%) compared to blood (≈10%). (**Fig. 4c**). CD8^+^ and CD4^+^ T cell subsets confirmed distinct functional states based on tissue type, with blood having higher levels of naïve and central memory markers (CD62L and CD27), and plaque having higher levels of the activation markers HLA-DR in both lineages and of CD38 exclusively in CD8^+^ T cells (**Extended Fig. 5a-e**). Plaque macrophages accounted for about 16% of CD45^+^ cells-(**Extended Fig. 5f, g**). Similar to our CYTOF results, we identified 2 macrophage subsets (CD64^+^HLADR^+^CD206^hi^ and CD64^+^HLADR^+^CD206^lo^) based on the varied expression of CD206, a known marker of alternatively activated M2 phenotype, which accounted for ~47% of plaque macrophages (**Extended Fig. 5h, i**).

**Figure 4.**
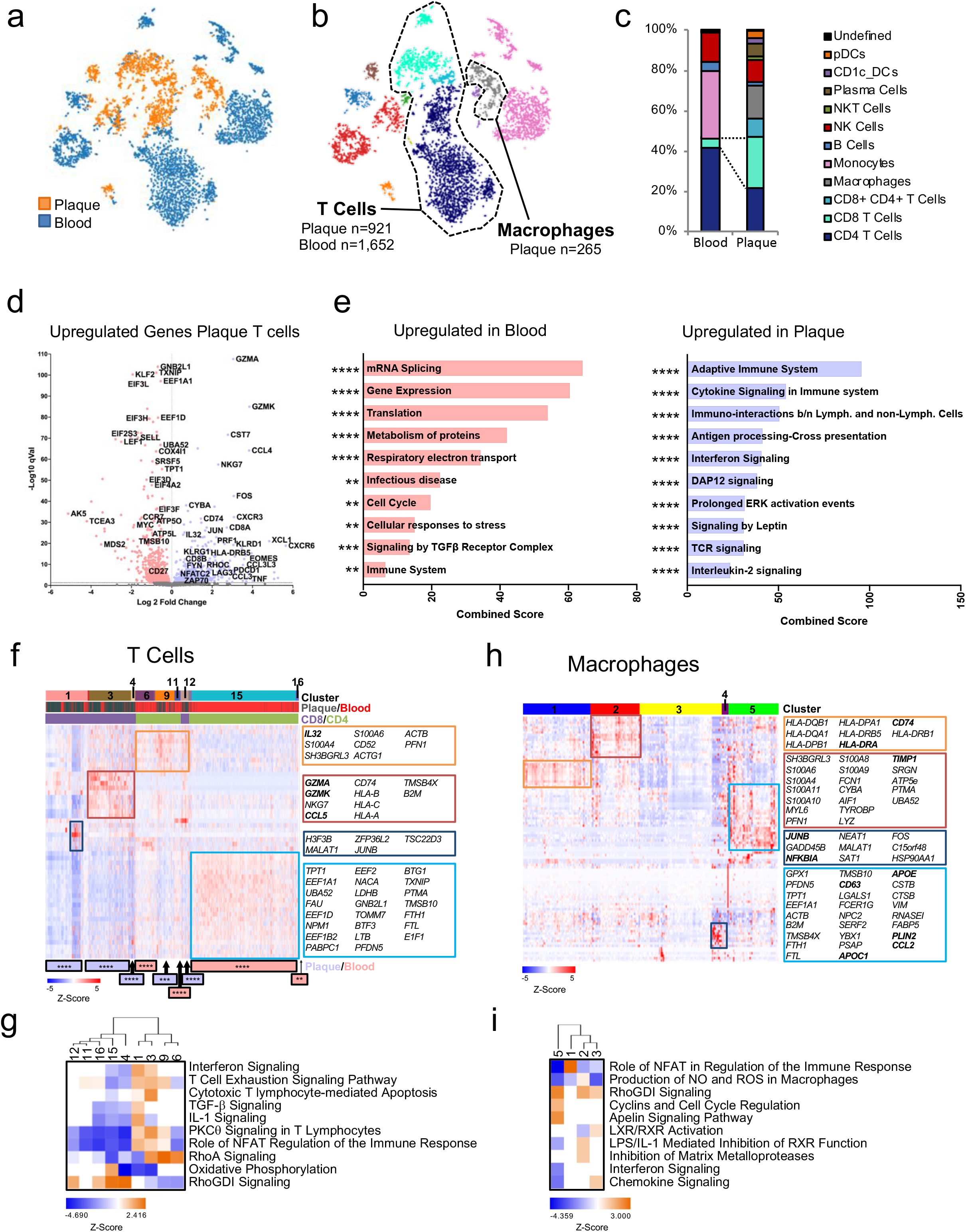
Simultaneous Epitope and Transcriptomic Analysis of Paired Atherosclerotic Plaque and Blood: CITE-seq Analysis. (a, b) viSNE plots of cells clustered from the Antibody Derived Tag (ADT) expression overlaid by (a) tissue, or (b) immune cell type. (c) Frequency of immune cells in blood and plaque. (d-i) Analysis of the gene expression data (GEX) from T cells (plaque n=921; blood n=1,652) and macrophages (n=265). (d) Volcano plot of the top 5000 DEGs in T cells in plaque (purple) or blood (pink). (e) Pathway analysis of T cell gene upregulated in blood or plaque. Bars indicate the combined score [p value (Fisher’s exact test) multiplied by the Z-score of the deviation from the expected rank] from *Enrichr*. (f) Heatmap of hierarchically clustered top 50 variable genes across all T cells in plaque and blood. Rows: z-scored gene expression values; columns: individual cell barcode. Identified cell clusters are numbered and color-coded. Bottom: cluster enrichment in blood (pink) or plaque (purple) and correspondent p values. Boxes list key genes in correspondent clusters. p values,**p<0.01, ***p<0.0001, ****p<0.0001. (g, i) Canonical signaling pathway analysis of the top 5000 DEGs per cluster corresponding to T cells (g) and Macrophages (i).

### GEX analysis of T cells in blood and atherosclerotic plaques

Next, we analyzed the single-cell transcriptional data of T cells in plaque and blood (**Fig. 4d**). Plaque T cells displayed gene expression signatures associated with T cell activation (*NFATC2, FYN, ZAP70*), cytotoxicity (*GZMA, GZMK*), and T cell exhaustion (*EOMES, PDCD1, LAG3*). These processes were reflected by signaling pathway analysis that highlighted T cell effector functions (Interferon and DAP12 Signaling^60,61^), proliferation and survival (IL-2 Signaling^62^), metabolic changes (Leptin Signaling^63,64^), and Cytokine Signaling pathways (**Fig. 4e**). In contrast, T cells in blood presented genes associated with cytokine inhibition (*KLF2*)^65^, RNA synthesis (*EEF1A1, EIF3L, TCEA3*) and metabolic reprogramming of circulating T cells (*TXNIP*) (**Fig. 4d**). These cells upregulated TGF-β signaling, RNA processing and splicing (*HNRNPM, SRSF5, RBMX*), and metabolism (*EIF4A2, PABC1, UBA52*) pathways, processes generally inhibiting T cell functions^66^ (**Fig. 4e**). Collectively, suggesting a resting phenotype in blood.

Unbiased hierarchical clustering of individual T cells identified 16 clusters with distinct enrichments in plaque or blood (**Fig. 4f**). Plaque-enriched clusters mostly corresponded to CD8^+^ T cells (1, 3, 4, and 12), and one CD4^+^ T cell cluster (9), confirming our CyTOF observations that plaques are enriched in CD8^+^ T cells and are composed of functionally heterogeneous subsets. Blood-enriched clusters mostly corresponded to CD4^+^ T cells (6, 11,15, 16). The clusters fell into two main functional categories, either activated or resting subsets (**Fig. 4g**). Blood-enriched (11, 15, 16) and plaque-enriched (4, 12) clusters presented a resting phenotype consistent with upregulated Rho GDP-dissociation inhibitor (RhoDGI) signaling, which antagonizes processes involved in T cell activation^67^. Blood-enriched Cluster 15 also displayed genes associated with RNA processing and Oxidative Phosphorylation (*ATP5MG, COX4I1, UQCRB*), suggesting their involvement in regulating T cell functions^66^. Conversely, plaque-enriched clusters 1, 3, 9 and blood-enriched cluster 6 presented distinct T cell activation states based on the differential representation of several canonical pathways. For example, RhoA signaling^68^ was upregulated in clusters 3, 6, and 9, and PCKθ signaling^69,70^ was upregulated in clusters 1, 3 and 9. Plaque-specific clusters 1, 3, and 9 uniquely upregulated signaling associated with activation-induced T cell exhaustion, including PKCθ signaling, required for TCR-induced T cell activation^71^, and NFAT signaling^72,73^. Overall, these data confirm our CyTOF observations that in blood, T cells display a quiescent state, while in plaques, they exhibit distinct degrees of activation which overlaps with exhaustion, suggesting the stepwise and progressive loss of T-cell functions in response to chronic, persistent inflammation^55–57,74^. A sub analysis of individual CD4^+^ and CD8^+^ T cells across blood and plaque is available as Supplementary Information (**Supplementary Fig. 5**)

### GEX analysis of macrophages in atherosclerotic plaques

The transcriptional profile of lesional macrophages was analyzed using hierarchical clustering, that identifed 5 distinct clusters. This method revealed a greater functional heterogeneity compared to the two subsets detected in our CyTOF and CITE-seq proteomic analysis (**Fig. 4h**). Signaling pathway analysis revealed that clusters 1, 2 and 3, were more activated and pro-inflammatory than cluster 5, which presented a foam cell transcriptional signature (**Fig. 4h and i**). Cluster 1 expressed genes involved in macrophage migration^75^ (MHCII genes [i.e. *HLA-DRA*] and *CD74*) and demonstrated upregulation of NFAT signaling, which dampens innate inflammatory responses through cytokine inhibition^76^. Cluster 2 was highly inflammatory, expressing genes involved in inflammatory responses (*CYBA, LYZ, S100A9, AIF1, S100A8*), toll-like receptor (TLR) binding (*S100A9, S100A8*), and oxidoreductase activities (*CYBA*). Interestingly, cluster 2 upregulated genes involved in matrix metalloprotease (MMP) suppression (*TIMP1*), a process that regulates extracellular matrix (ECM) degradation^77^. This data suggests that these cells might contribute to plaque stabilization. Cluster 3 showed reduced gene expression compared to other clusters. However, these cells uniquely upregulated genes involved in pro-inflammatory responses (*JUNB*, *NFKBIA*)^78^ and highly expressed *MALAT1*, a long non-coding gene implicated in M2 macrophage polarization^46,79^ and foam cell formation^80^. Cluster 3 upregulated liver X receptor (LXR) and retinoid X receptor (RXR) activation signaling, processes that modulate reverse cholesterol transport and efflux^81^, suggesting these were precursory foam cells. Finally, cluster 5 upregulated genes involved in cholesterol uptake and metabolism (*PSAP*, *APOC1*, *APOE*) and lipid accumulation (*PLIN2*)^82^, and may represent a foam cell subset functionally distinct from cluster 3. These macrophages showed increased RhoGDI and Apelin signaling, and reduced NO/ROS production and pro-inflammatory cytokine signaling (e.g. IL-1, IFN). With the exception of the pro-atherogenic chemokine *CCL2*^83^, this data suggested an anti-inflammatory nature for macrophages of cluster 5.

### Discrete Immune Dysregulations Linked to Clinical Cerebrovascular Outcomes

#### Single-Cell Proteomic Analysis of ASYM and SYM plaques

To identify the plaque-specific immune perturbations associated with ipsilateral acute ischemic cerebrovascular events, we stratified the profiles of SYM and ASYM patients from the T cell MC analysis (Cohort 2, **Fig. 3, 5a**). We first analyzed MC frequency and found that SYM patients had a greater frequency of MC12 (CD4^+^ EM T cells) in the culprit plaque (≈32%, SYM vs. ≈20%, ASYM) (**Fig. 5b-d**, **and Supplementary Fig. 6a**). Remarkably, these changes were unrelated to the plaque type V or VI seen in cohort 2 based on the pathological criteria of the AHA^84^, the current gold standard to identify vulnerable/complicated atherosclerotic lesions (**Fig. 5e-g, See Supplementary Information, Supplementary Fig. 2f-h**). Surface marker expression of MC 12 did not identify significant phenotypic differences in MC 12 of SYM and ASYM patients (**see Supplementary Information, Supplementary Fig. 6b**).

**Figure 5.**
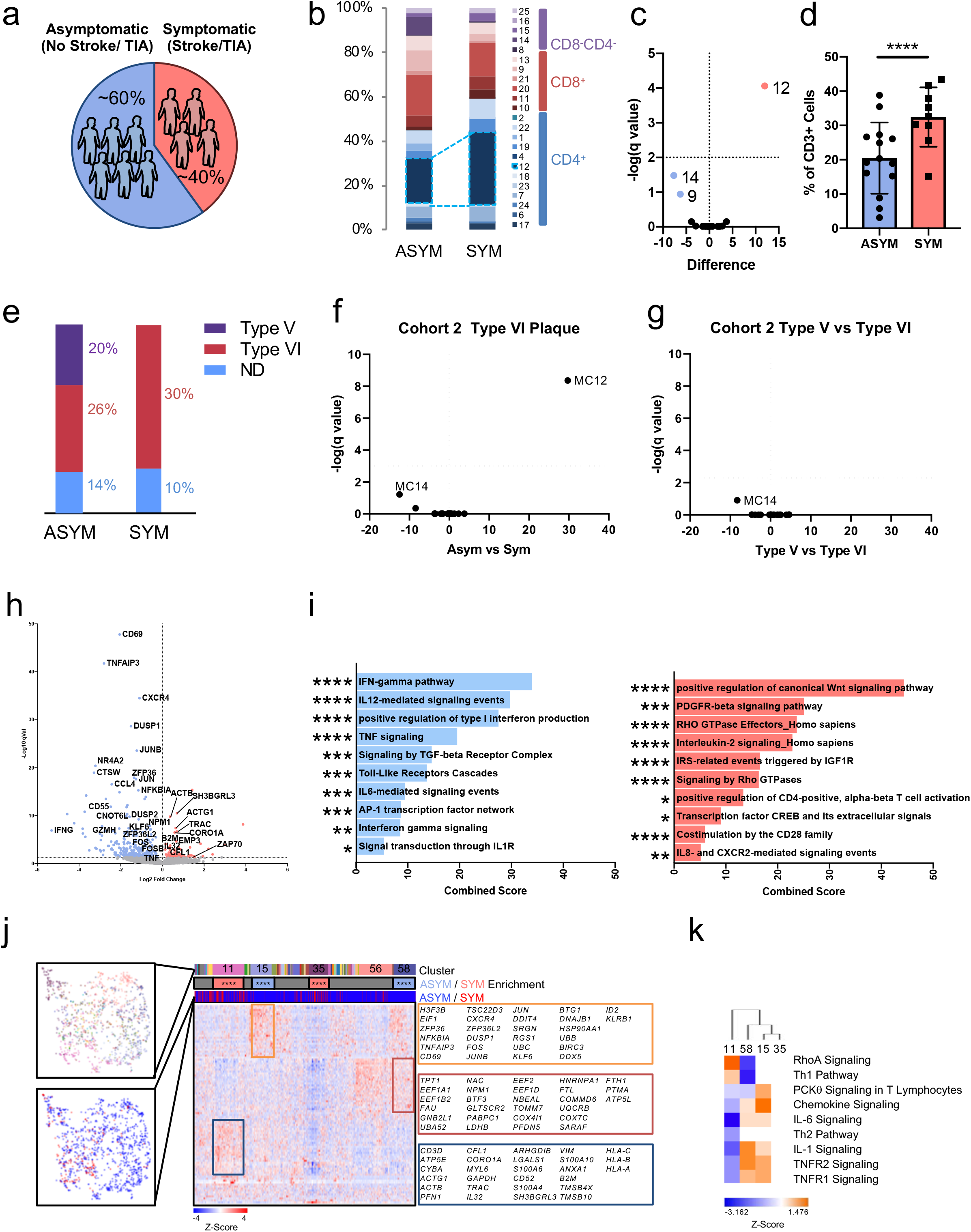
CD4^+^ T Cell Dysregulations Associated with Cerebrovascular Events. (a) Patients by clinical association. (b) T cell MC frequencies in plaque from the validation cohort stratified by asymptomatic (ASYM, n=14) and symptomatic (SYM, n=9) patients. (c) Volcano plot of T cell MCs enriched in SYM (red), or ASYM (blue) patients. P values were calculated using multiple t-Test with an FDR=0.5% (Benjamini, Krieger and Yekuteli method) (d) Frequency of MC12 per each patient. ****q<0.00001. Values are mean ± SD. (e) AHA plaque type classification in SYM and ASYM plaques of cohort 2 (n=23). (f) Assessment of T cell MCs per (f) clinical phenotype in type VI plaques from either SYM and ASYM patients, or (g) per plaque type (type VI vs. type IV) in all 23 patients. (h-k) Single-cell gene expression data of CD4^+^ T cells (n=1,200). (h) Top 5000 Differentially expressed genes across plaques of SYM and ASYM patients. (i). Signaling pathway analysis of the upregulated genes of CD4^+^ T cells from SYM or ASYM. (j) Unbiased hierarchical clustering of the top 100 variable genes across all CD4 T Cells. Column categories from top to bottom indicate cluster number, cell cluster enrichment in SYM/ ASYM, and cells from SYM (red) and ASYM (blue) subjects. (k) Canonical signaling pathway analysis of SYM and ASYM enriched clusters.

#### Single-Cell Transcriptomic Analysis of ASYM and SYM plaques

To further investigate discrete adaptive immune alterations and identify functional dysregulations of macrophages associated with clinical cerebrovascular outcomes, we performed a scRNA-seq analysis of T cells and macrophages in plaques of SYM and ASYM patients.

#### CD4^+^ T cells Transcriptional Alterations in ASYM and SYM Plaques

Gene set enrichment analysis of differentially expressed genes (DEGs) in CD4^+^ T cells from SYM vs. ASYM plaques (**Fig. 5h-i**) highlighted a transcriptional profile consistent with stronger activation and effector functions in ASYM patients. ASYM plaques upregulated genes involved in IL-12 signaling, type I interferon production, and increased *IFNG* expression, which collectively indicate activated effector functions of T cells^61^. Interestingly, these cells also exclusively upregulated IL-1 and IL-6 signaling pathways (**Fig. 5i**). Because IL-1 acts as a licensing signal that stabilizes cytokine transcripts and enables effector functions^85^ and IL-6 promotes T cell survival^86^, these pathways may enhance effector functions of CD4^+^ T cells in ASYM plaques. Conversely, SYM CD4^+^ T cells explicitly lacked IL-1 and IL-6 signaling pathways, which may indicate that therapeutics that target these pathways^33^ may be less effective in targeting plaque specific immune responses following acute ischemic events. Furthermore, CD4^+^ T cells from SYM plaques displayed distinct signaling pathways involved in T cell migration (RhoGTPase)^68^, T cell activation (PDGFR-β)^87^, T cell differentiation (Wnt^88^, IL-2^62^) and T cell homeostasis (CD28).

Unbiased hierarchical clustering of individual CD4^+^ T cells across all plaques identified 4 main clusters with distinct enrichments between ASYM (clusters 15 and 58) and SYM (clusters 11 and 35) plaques (**Fig. 5j**). ASYM-enriched clusters shared many of the same pathways highlighting a similar activated cell phenotype (**Fig. 5k**). Conversely, SYM-enriched cluster 11 was activated and proinflammatory, as indicated by Th1 pathway and RhoA signaling, while SYM-enriched cluster 35 presented an overall downregulation of gene expression.

#### CD8^+^ T cells Transcriptional Alterations in ASYM and SYM Plaques

Gene expression analysis of CD8^+^ T cells in ASYM and SYM plaques identified a transcriptional profile (**Fig. 6a**) which partially overlapped with that of CD4^+^ T cells (**Fig. 5h**). CD8^+^ T cells in ASYM plaques upregulated activation genes (*CD69, NFKBIA, IFNG*, and *TNF*) and signaling pathways (type I interferon signaling, TNF signaling, IL-6 signaling) (**Fig. 6b**). These patterns of upregulation were shared between T cell lineages, suggesting a similar activation state. However, upregulation of unique genes (*CXCR4, GZMB, CCL4L2*) and signaling pathways (granzyme, TGFβ) indicated specific chemotactic and cytotoxic functions of these CD8^+^ T cells in ASYM plaques. In contrast, SYM CD8^+^ T cells were largely characterized by a distinctive set of genes (*RAC2, CCL5, CYBA, GZMK, LIMD2*) (**Fig. 6a**) and signaling pathways (TCR, CXCR2, cellular senescence, and IFNγ) (**Fig. 6b, c**). Hierarchical clustering of all CD8^+^ T cells identified 5 clusters with distinct enrichments between ASYM (clusters 3 and 4) and SYM (clusters 1, 5 and 7) plaques (**Fig. 6d**). Signaling pathway analysis (**Fig. 6e**) revealed that ASYM-enriched cluster 4 was characterized by inflammatory (IL-6, chemokine signaling) and regulatory pathways related to T cell activation and migration (i.e. RAC signaling, TNFR1/2 signaling, TGFβ signaling). This suggests that activation signaling is tightly regulated in this cluster. SYM-enriched cluster 1 and ASYM-enriched cluster 3 shared a similar, activated functional state characterized by RhoA, Integrin, IL-8 and NFAT signaling pathways (**Fig. 6e**). However, cluster 1 was distinctive in the co-activation of Interferon and T cell exhaustion signaling (**Fig. 6e**), suggesting that these highly activated cells initiate a parallel exhaustion reprogramming. Because T cell exhaustion is characterized by the gradual and continuous depletion of effector functions in response to chronic, non-resolving inflammation^55–57,74,89^, the identification of concomitant activation and exhaustion signaling suggests that these activated cells initiate exhaustion reprogramming.

**Figure 6.**
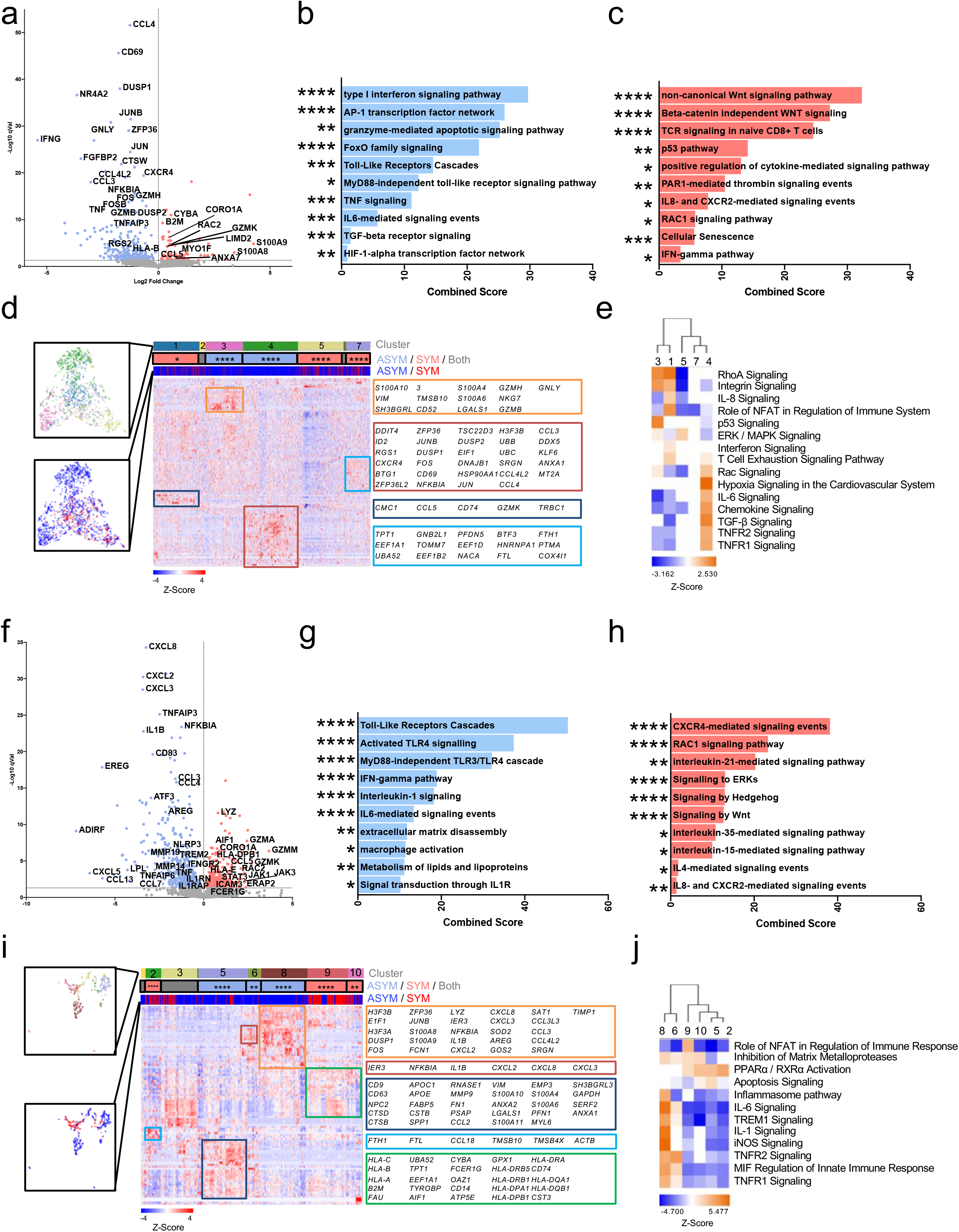
Transcriptional Dysregulations Associated with Cerebrovascular Events in CD8^+^ T Cells and Macrophages. Differential gene expression of the top 5000 differentially expressed genes (DEGs) across SYM and ASYM plaques (a) CD8^+^ T Cells (n=1,747) and (f) macrophages (n=747) was calculated using the Welch’s T-test using UMI normalized gene expression data. *Enrichr* signaling pathway analysis of CD8^+^ T cells (b, c) and macrophages (g, h) on the top 5000 DEGs between SYM or ASYM patients. Hierarchically clustered cells based on the Z-scored expression of the top 100 variable genes across all CD8^+^ T cells (d) and macrophages (i). Canonical pathway analysis of SYM and ASYM enriched clusters in CD8^+^ T cells (e) or Macrophages (j).

#### Macrophage Transcriptional Alterations in ASYM and SYM Plaques

Similar to the T cell compartment, ASYM macrophages were more activated, pro-inflammatory, and displayed enhanced foam cell functions compared to SYM macrophages. ASYM macrophages highly expressed a series of pro-inflammatory chemokines (**Fig. 6f**) corresponding to signaling pathways including IL-1 and IL-6 (**Fig 6g**). In fact, IL-1 regulation was highly represented in ASYM plaques, which upregulated genes *IL1B* and *IL1RAP*, a component of the IL-1 receptor complex, and *NFKBIA* and *NLRP3* mediators of IL-1 β production^90^ (**Fig 6f**). Interestingly, inhibitory IL-1 decoy receptor (*IL1RN*), was co-expressed, indicating tight self-regulatory mechanisms for IL-1β signaling in macrophages of ASYM plaques (**Fig 6f**). Additionally, these macrophages upregulated genes involved in lipid metabolism and lipoproteins implicated in foam cell formation (e.g. *LPL*), a hallmark of all stages of atherosclerosis^1,21,59^.

In SYM plaques, macrophages upregulated cytolytic genes *GZMA, GZMK, GZMM* and *LYZ*, as well as the proatherogenic chemokine *CCL5* that has been linked to an unstable plaque phenotype^91,92^(**Fig. 6f**). Signaling pathway analysis revealed heterogeneous populations of inflammatory, reparative, pro-angiogenic, and anti-fibrotic macrophages (**Fig. 6h**). Top pathways included CXCR4 signaling, involved in cell migration and TLR2 signaling^93,94^, which indicates pro-inflammatory functions of these cells. We also identified pathways involved in podosome formation (RAC1^95,96^), and phagocytosis (RAC1^97^ and IL-21^98^ signaling), both processes that promote reparative functions in macrophages. We also identified pathways regulating macrophage survival and wound healing (IL-35 signaling^99^) (**Fig. 6h**). Furthermore, we identified pro-angiogenic IL-8 and CXCR2 signaling, which have been implicated in plaque instability^100–103^, and the Hedgehog and Wnt signaling, both involved in anti-fibrotic responses which may contribute to plaque destabilization^104–106^ in SYM macrophages.

Unbiased hierarchical clustering revealed heterogeneous and highly specialized macrophage subsets enriched in either ASYM plaques (clusters 5, 6 and 8) or SYM plaques (clusters 2, 9 and 10) (**Fig. 6i**). Consistent with our initial analysis, ASYM macrophages were either phenotypically activated and pro-inflammatory (cluster 6 and 8) or demonstrated a foam cell profile (cluster 5) (**Fig. 6i, j**). Macrophages from cluster 8 expressed proinflammatory genes *IL1B*, *CXCL2*, *CXCL3* and *CXCL8*, and upregulated activation pathways (inflammasome pathway, IL-1, IL-6, iNOS, MIF and TREM1 signaling) (**Fig. 6i and j**). Cluster 5 uniquely upregulated genes involved in lipid uptake and metabolism (*APOC1*, *APOE*, *FABP5*) and cholesterol transportation (*NPC2*) suggesting that these macrophages were foam cells (**Fig. 6i**). Cells in this cluster also upregulated apoptosis signaling. In contrast, clusters from SYM plaques 2, 9 and 10 displayed fewer proinflammatory functions (i.e. NFAT signaling) and stronger signaling associated with alternatively activated phenotypes linked to plaque instability (**Fig. 6j**). Clusters 9 and 10 shared similar phenotypes including PPAR α/RXRα signaling that has been reported to drive macrophage polarization towards the reparative M2 phenotype^107–111^. Cluster 9 was distinct based on the activation of NFAT signaling, which is a shared signaling to that of macrophages in rheumatoid arthritis^112^ and inflammatory bowel disease^113^, conditions that are at high risk of CV disease^114,115^. Cluster 2 was uniquely characterized by the upregulation of genes responsible for iron metabolism (*FTH1*) and iron storage (*FTL*). This cluster may represent alternatively activated pro-inflammatory non-foamy macrophages with proatherogenic potential, which are known to be involved in the clearance of iron derived from hemoglobin deposited in areas of intraplaque hemorrhage^47^. This phenomenon has been associated with plaque progression and features of plaque instability and is typically seen in AHA type VI complicated plaques^84,116,117^, the same type that is exhibited by all SYM patients in our cohort.

### Cell-to-Cell Communications in Atherosclerotic Plaques Associated with Cerebrovascular Events

We next utilized an established computational approach^118^ to infer cellular communication mechanisms^119,120^ by quantifying the potential ligand-receptor interactions between cell types based on their single-cell gene expression profiles. Using this approach^118^, we identified cell-to-cell communication mechanisms that may contribute to the distinct functional state of T cell and macrophage in atherosclerotic plaques of ASYM and SYM patients (**Fig. 7 and Extended Fig. 6, see Supplementary Information, and Supplementary Figs. 7-8**).

**Figure 7.**
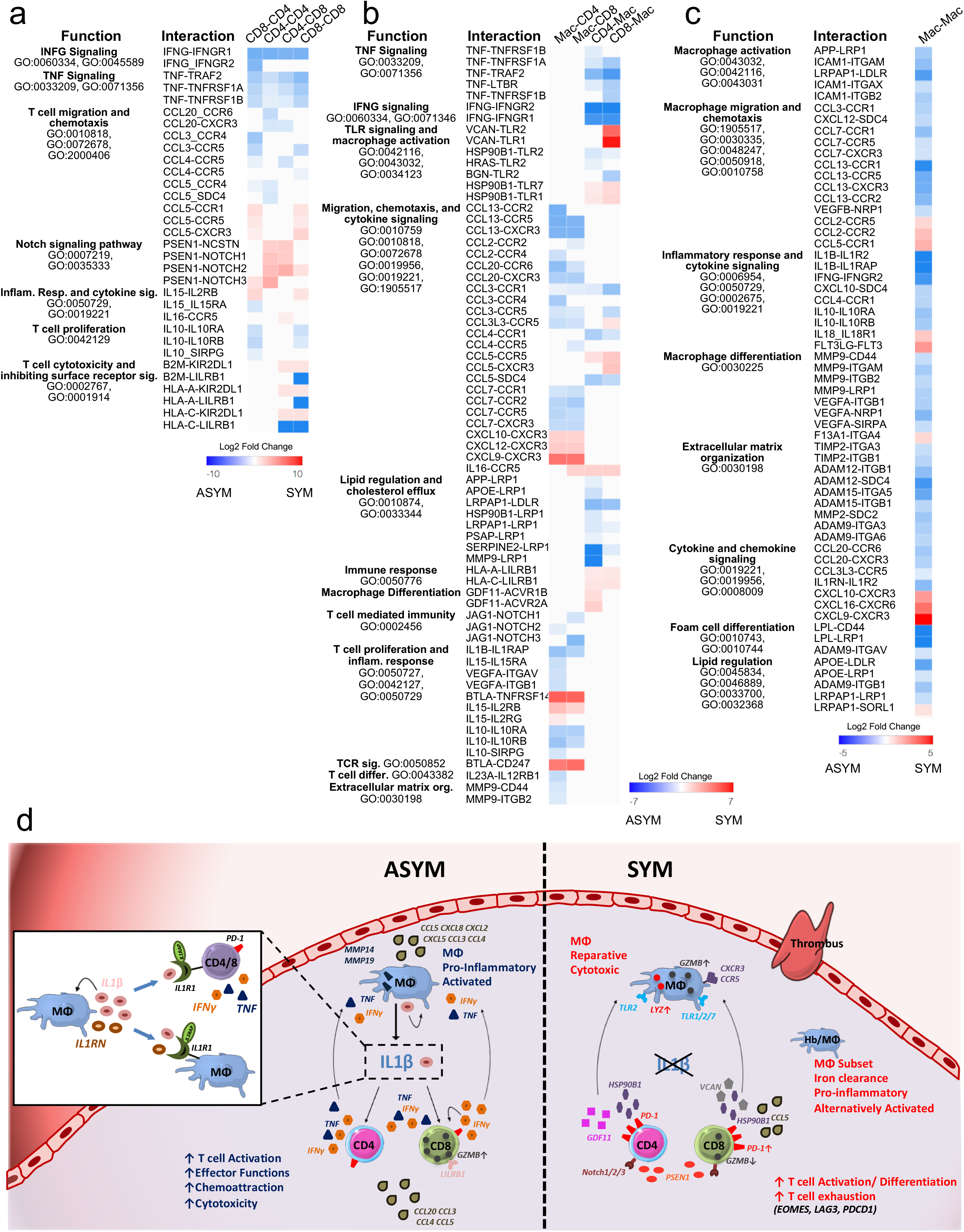
Cell-Cell Interactions Associated with Cerebrovascular Events. Ligand-Receptor interactions involving the top 30 SYM/ASYM differentially regulated ligands based on absolute value sum of log2 fold change across ligands and visualized using *Clustergrammer*. (a) T cell-T cell, (b) T cell-Macrophage and Macrophage-T cell, and (c) Macrophage-Macrophage interactions are presented as (ligand-receptor). Heatmaps represent the Log2 fold change of the interaction score between SYM and ASYM. Log2 foldchange values were clipped at +/-10. (d) Visual summary of the key innate and adaptive immune functional alterations and main differences in cell-cell communications seen in plaques from ASYM and SYM patients.

#### T cell-T cell Interactions in ASYM and SYM plaques

In ASYM plaques, we identified potential ligand-receptor interactions between CD4^+^ and CD8^+^ T cells that could sustain the activated, pro-inflammatory phenotype, seen in both T cell lineages, for example through TNF and IFNγ signaling (**Fig. 7a**; **Extended Fig.6a, b; Supplementary Figs. 7a,b, 8a,b**). Additionally, we found ligand-receptor interactions involved lymphocyte chemotaxis and migration^121^, including cytokines and chemokine ligands that interact with receptors on both CD4^+^ and CD8^+^ T cells. In SYM plaques, we identified distinct interactions known to be involved in key functional states like T cell activation, differentiation, and exhaustion. Interestingly, both CD4^+^ and CD8^+^ T cells expressed *PSEN1*, a gamma secretase component known to cleave Notch receptors (*NOTCH1/2/3*)^122^ that were expressed by both T cell linages in plaque. This signaling is involved in T cell function and may contribute to the observed T cell activation^123^, differentiation^124^, PD-1 expression and regulation of cytolytic functions^125^ in our plaques. Our analysis also revealed that T cells expressing the same ligands could interact through distinct receptor pairs on CD4^+^ and CD8^+^ T cells in SYM and ASYM plaques. For instance, *IL-15* expressed by CD8^+^ T cells could modulate both CD4^+^ and CD8^+^ T cell functions, but through different receptors expressed by these cells in ASYM and SYM plaques. In ASYM plaques, we identified *IL-15-IL15RA* on CD4^+^ T cells, an interaction involved in cell proliferation, cell migration, and inhibition of apoptosis^126^. On the other hand, in SYM plaques *IL-15-IL2RB* on both CD4^+^ and CD8^+^ T cells could regulate Th1 reprogramming^127^ and differentiation^128^, two functional phenotypes that emerged in SYM plaques by from our CyTOF and scRNA-seq analyses.

#### T Cell-Macrophage Interactions in ASYM and SYM plaques

Identified ligand-receptor interactions between T cells and macrophages in ASYM plaques highlighted unique adaptive signals involved in macrophage activation, migration, regulation of lipids and foam cell formation, as well as innate immunomodulatory signaling (**Fig. 7b**; **Extended Fig. 6a, b, Supplementary Fig. 7a-c**). Specifically, T cell-macrophage communications involved the activating and pro-inflammatory interactions *TNF*-*TNFRSF1A*/*1B* and *IFNG*-*IFNGR1*/*2*. T cells expressed ligands (*HSP90B1, BGN, HRAS*) that were predicted to stimulate TLR2 signaling on macrophages, a key pathway that regulates the pro-inflammatory status of macrophages in human atherosclerosis^129^. Consistent with the specific identification of foam cells in ASYM plaque from our scRNA-seq analysis, we found T cell-macrophage interactions in ASYM plaques known to regulate lipid accumulation and foam cell formation though the expression of multiple ligands that bind to the proatherogenic LDL Receptor-Related Protein (*LRP1*) on macrophages^130^. In SYM plaques, T cells might also stimulate macrophage activation through unique TLR signaling cascades by which ligands (*VCAN, HSP90B1*) bind to TLRs (*TLR1/2/7*). Additionally, in SYM plaques we identified ligand-receptor interactions between T cells and macrophages involved in cell survival (*HLA-A/C-LILRB1 and B2M-LILRB1*)^131^ and pro-atherogenic functions linked to plaque vulnerability (*CCL5-CCR5*)^91,92,132^. Interestingly, T cell expressed ligands that may promote the reparative macrophage polarization detected in these plaques by scRNA-seq analysis, through the expression of GDF11, a TGFβ superfamily member which bound to *ACVR2A/1B* on macrophages^133^.

In ASYM plaques, we identified macrophage ligands interacting with CD4^+^ and CD8^+^ T cell receptors that could promote T cell activation (*JAG1*-*NOTCH1/3*)^134^, and T cell recruitment through the expression of distinct T cell chemokines. Importantly, ASYM macrophages exclusively expressed *IL1B* that was predicted to bind to the IL-1 receptor accessory protein *IL1RAP* expressed by CD4^+^ and CD8^+^ T cells, an interaction that may enhance effector functions of T cells infiltrating ASYM plaques^85^. This data are in line with our findings from scRNA-seq analysis that IL-1 β signaling is exclusive to ASYM plaques. On the other hand, in SYM plaques, we identified fewer ligand-receptor interactions between macrophage and T cells. Among those, macrophages modulated T cell activation through the interactions between *IL15-IL2RB/G* and *BTLA*-*CD247*, the latter of which is a component of the TCR-CD3 complex required for T cell activation^135^. Macrophages could also promote T cell recruitment^136^ into SYM plaques through their of chemokines-receptor interactions (**Fig. 7b**; **Extended Fig. 6a, b; Supplementary Fig. 7a-c**).

#### Macrophage-Macrophage Interactions in ASYM and SYM Plaques

We detected more diverse and distinctive communication patterns in autocrine macrophage interactions. Specifically, identified macrophage-macrophage interactions in ASYM plaques could induce self-activation and pro-inflammatory interactions via *ICAM1* binding to *ITGAX*, *ITGB2* and *ITGAM*, self-recruitment via *CCL7* and *CCL13* binding to *CCR1*, *CCR2*, *CCR5* and *CXCR*3, and activation of inflammatory IL-1 signaling via *IL1B* binding to *IL1R2 and IL1RAP*. Also macrophages could stimulate MMP activity and angiogenesis as seen with *MMP9*, *ADAM9*, *ADAM12*, *ADAM15* and *VEGFA* signaling through receptors *ITGB1*, *ITGAV*, *ITGA3*, *ITGA5*, *ITGA6*, and *NRP1*, indicating a strong influence of macrophages in driving ASYM plaque development and progression. In SYM plaques, lesional macrophages could activate antigen processing and presentation through *HLA-C* upregulation and might promote a proinflammatory state via *IL18* signaling through *IL18R1*, and *FLT3 to FLT3LG*. Interactions were also observed between chemoattractants *CCL2*, *CCL5*, *CXCL9*, *CXCL10* and *IL16* that bind *CXCR3*, *CCR2* and *CCR5* receptors. Additionally, our analysis showed that SYM macrophages may be involved self-maintaining the reparative M2-like phenotype seen in these plaques through the expression of the M2 marker *F13A1*^137^ (**Fig. 7c**; **Extended Fig. 6a, b; Supplementary Fig. 8c**).

## DISCUSSION

In this study, we provide a first in depth single-cell CyTOF and CITE-seq immune mapping of plaques and paired blood in patients with atherosclerosis. Importantly, unbiased high-dimensional single-cell CyTOF and scRNA-seq analyses uncovered new specific innate and adaptive immune dysregulations at the atherosclerotic site associated with clinical cerebrovascular events.

Understanding the contribution of the immune system to the development of human atherosclerosis has been a daunting task. Recently, human plaque immune composition of has been inferred using deconvolution methods on bulk RNA-seq data from human atherosclerotic tissue and proof-of-concept CyTOF analysis of an individual plaque^38^. A main limitation of this approach is that bulk RNA-seq averages of gene expression patterns in the whole tissue limiting the resolution to identify biologically relevant functional variations of individual cells. Single-cell technological advances allow precise measurements of individual cell phenotypical and functional variations^138–141^ that have the potential of changing our conceptual understanding of human vascular biology and disease state.

Our study is the first and the largest to integrate single-cell CyTOF, CITE-seq and scRNA-seq to study the immune composition of human plaques and to identify the plaque specific immune dysregulations associated with clinical cerebrovascular events at unprecedented scale and resolution. We designed unbiased CyTOF and CITE-seq studies to directly compare immune cell repertoires from paired plaque and blood from the same patient. These analyses underscore unique differences and highly specialized functions of plaque T cells. Not only were T cells prominent in atherosclerotic lesions, but they were overall more activated, differentiated, and exhausted compared to their blood counterparts. In particular, we identified an enrichment of 4 specific effector memory T cell subsets in plaques that presented distinct activation and differentiation patterns indicating a functional heterogeneity of the T cell immune compartment. Interestingly, one of these subsets corresponded to a CD103^+^ tissue resident memory (TRM) CD8^+^ T cells, that have previously been described in other non-lymphoid human tissues^48^, but never in the atherosclerotic arterial wall. A key finding of our CyTOF analysis was that plaque T cells expressed high levels of PD-1, a marker of T cell exhaustion reprogramming that is upregulated upon T cell activation^68–71^, that was confirmed at the transcriptional level by our CITE-seq analysis. We found that plaque T cells expressed a correspondent exhaustion transcriptional signature (i.e. *EOMES, LAG3*) that parallel that of exhausted T cells in the tumor microenvironment^56^. CITE-seq analysis also revealed the co-existence of exhausted and activated T cell subsets within the same plaque, suggesting that highly activated T cells may initiate an exhaustion reprogramming, the stepwise and progressive loss of T cell functions, possibly sustained by chronic unresolving plaque inflammation^55–57,74,89^. CyTOF analysis on plaque-derived immune cells also identified two main subsets of macrophages, resembling the classically-activated M1 and alternatively-activated M2 phenotypes^46,47^. Interestingly, plaque macrophages were more resolved at the single-cell transcriptional level in our CITE-seq analysis, indicating a diverse functional heterogeneity of these cells in plaque. We found distinct macrophage subsets displaying activated and pro-inflammatory functional states, inflammatory macrophages expressing genes involved in lipid metabolism, a subset distinctively resembling foam cell functions, and macrophages with unique anti-inflammatory transcriptional signatures. These data highlight the functional specialization and adaptation of lesional macrophages to the pro-inflammatory, lipid rich human plaque microenvironment that is not accurately reflected by the M1 and M2-phenotypes resolved by canonical surface protein marker analysis^17,21,46^.

Our combined analyses of CyTOF and scRNA-seq data underscored new innate and adaptive immune dysregulations and alterations of key cell-cell communications in the plaque microenvironment associated with clinical cardiovascular outcomes. We found that ipsilateral plaques of symptomatic patients had significant adaptive dysregulations consistent with an expansion of activated effector memory CD4^+^ T cell subset. Interestingly, this change was not related to either plaque pathology according to the AHA classification^84^ or clinical characteristics.

Single-cell transcriptomic analysis of plaque T cells highlighted more heterogeneous distinctive alterations of both CD4^+^ and CD8^+^ T cells associated with recent clinical events. Specifically, in ipsilateral plaques of symptomatic patients with recent stroke or TIA, both T cell lineages presented gene expression signatures largely consistent with activation, differentiation, and exhaustion. Conversely, T cells were overall activated in plaques of asymptomatic patients. Cell-cell interaction analysis showed that these functional states were tightly regulated by specific interactions with other T cells and macrophages. Previous murine studies have shown that PD-1 signaling, a major regulator of T-cell exhaustion, is required to modulate atherogenic responses of activated T cells in the arterial wall and its inhibition results in aggravated atherosclerosis^142^. Our data that T cell exhaustion is subset-specific and co-exists with activated T cells in ipsilateral human plaques from symptomatic patients, suggests an unforeseen and more complex role of exhaustion in human disease that may have important clinical implications. The fundamental discovery that PD-1 inhibitors reverse anergy of tumor T cells has fundamentally changed the therapeutic options and prognosis for patients with advanced malignancy^53,54,143^. Our data that exhausted T cells expressing high levels of PD-1 exists in atherosclerotic plaques suggest that off-target effects of PD-1 inhibitors may increase T cell activation. Considering that T cell activation aggravates atherosclerosis^21^, treatment with PD-1 inhibitors may have unforeseen consequences in cancer patients with underlying atherosclerotic CV disease. Additionally, the higher exhaustion seen in subsets of plaques of patients with recent cerebrovascular events, suggest that any possible off-target effect may have higher consequences in patients with recent history of stroke or TIA which are already at higher risk of recurrent ischemic events within 5 years^144^. Future cardio-oncology monitoring studies designed to investigate unrecognized cardiovascular side effects of increasingly used immune checkpoint inhibitors^143^ would be useful to test this hypothesis and help improve patient selection and cardiovascular monitoring in the clinical setting.

Similar to the plaque T cells, macrophages infiltrating plaques of asymptomatic patients presented an activated and pro-inflammatory phenotype. Conversely, lesional macrophages from ipsilateral plaques of patients with recent CV events displayed a transcriptional signatures more consistent with distinct pro-inflammatory and reparative functions, including a proatherogenic subsets of cell involved in iron metabolisms derived from the hemoglobin deposited in areas of intraplaque hemorrhage that characterize AHA type VI complicated plaques^47,84,116,117^, the same plaque type that is exhibited by all SYM patients in our cohort. We found that the specialized functions of lesional macrophages in atherosclerotic plaques from symptomatic and asymptomatic patients were self-regulated but also dependent on T cell ligand signals, indicating unique and highly coordinated cross-communications between innate and adaptive immune responses at the lesional site.

A key and unexpected finding of our study was the discovery that IL-1 signaling, which was targeted in the CANTOS to reduce the CV risk of high risk post-MI patients^33^, was differentially regulated in plaques from symptomatic and asymptomatic patients. Plaque macrophages of asymptomatic patients expressed higher levels of *IL1B*, and associated IL-1 signaling compared to symptomatic patients. Additionally, we found that IL-1 did not exclusively signal innate pro-inflammatory functions in macrophages but also in T cells, suggesting a key role at the intersection of innate and adaptive immune responses by sustaining effector functions of T cells in ASYM plaques. These data suggest that post-MI patients in the CANTOS may have been less susceptible in their response to IL-1β inhibition at the culprit plaque site, than high-risk patients without recent CV event.

In conclusion, by combining proteomics and transcriptomics single-cell analyses of atherosclerotic plaques and blood of the same patients provides a first immune atlas of human atherosclerosis and identified new specific innate and adaptive immune dysregulations at the atherosclerotic site associated with clinical cerebrovascular events.

Our results strongly support the concept that an accurate selection of patients is required to optimize treatment efficacy, and that new immunotherapies must be precisely tailored to discrete immune molecular and cellular defects in subsets of patients^32,33,145,146^.

## ACKNOWLEDGMENTS

We thank the Human Immune Monitoring Center (HIMC), in particular Oksana Mayovska, Victor Guo, Xiaochen Ivy Qin, Hui Emily Xie, Manishkumar Patel, Melanie Davila, Brian Lee, Shermineh Bradford, Laura Walker, and Kevin Tuballes. We thank Swathy Sajja and Peik Sean Chong for their coordination efforts. We are grateful to Dr. Alice Kamphorst for her critical review of this manuscript. We thank the Biorepository and Pathology Core of the Icahn School of Medicine at Mount Sinai. This work utilized mass cytometry instrumentation supported by NIH grant S10OD023547-01. This work was funded by NIH grants K23HL111339, R03HL135289. C.G. was also funded by NIH grants R21TR001739, and UH2TR002067 and partially supported by the American Heart Association (14SFRN20490315). D.F. is supported by the NIH grant 5T32HL007824-20. A.M. and Z.W. are supported by NIH grants U54-HL127624 (LINCS-DCIC) and U24-CA224260 (IDG-KMC).

## AUTHOR CONTRIBUTIONS

Conceptualization: C.G., M.M., and A.H.R.; methodology: C.G., A.H.R., E.D.A., N.F., A.M., S.G., D.M.F. J.L.M.B.; software: E.D.A., N.F., Z.W., A.M.; formal analysis: A.H.R., D.M.F., E.D.A., N.F., Z.W., A.M. investigations: D.M.F., A.H.R., A.C., L.A., N.K., R.R., R.S., S.K-S. C.W., R.S. C.H.; Resources: J.R.L., C.P., N.M., C.F., A.J.A, J.M., P.F. and A.M.; data curation: A.H.R., D.M.F, N.F., E.D.A., S.K-S, J.L.M.B.; Writing: C.G; revision and editing: C.G., A.H.R., D.M.F., M.M., S.K-S, C.W., C. H., R.S., N.F.; data visualization: C.G., D.M.F., N.F., A.H.R., A.C., and A.M; supervision: C.G., A.H.R., S.K-S and M.M.; project administration: C.G.; funding acquisition: C.G.

---

**Extended Figure 1.**
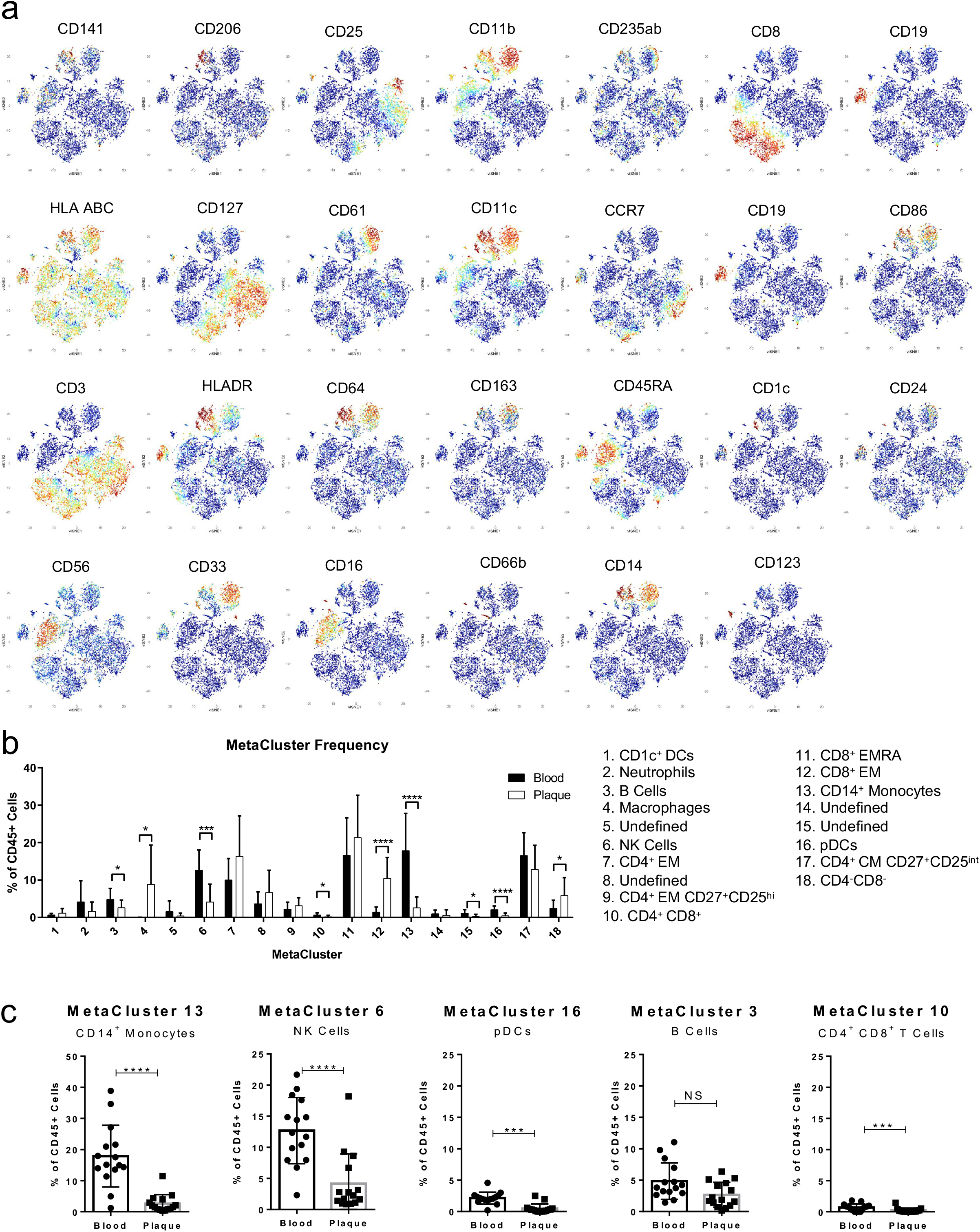
Definition of MetaClusters from the Discovery Cohort. (a) viSNE plots displaying the protein marker overlaid in spectral colors. (b) Bar plot of MC frequencies in blood (black) and plaque (white). Annotated MCs are shown on the right. Statistical significance was determined using unpaired t-test with multiple comparisons corrected by FDR=0.5%. *q<0.05, ***q<0.001, and ****q<0.0001, (c) Comparison of population frequency of MCs enriched in blood. ***p<0.001, ****p<0.0001 by Wilcoxon test. Values of all plots are mean ± SD.

**Extended Figure 2.**
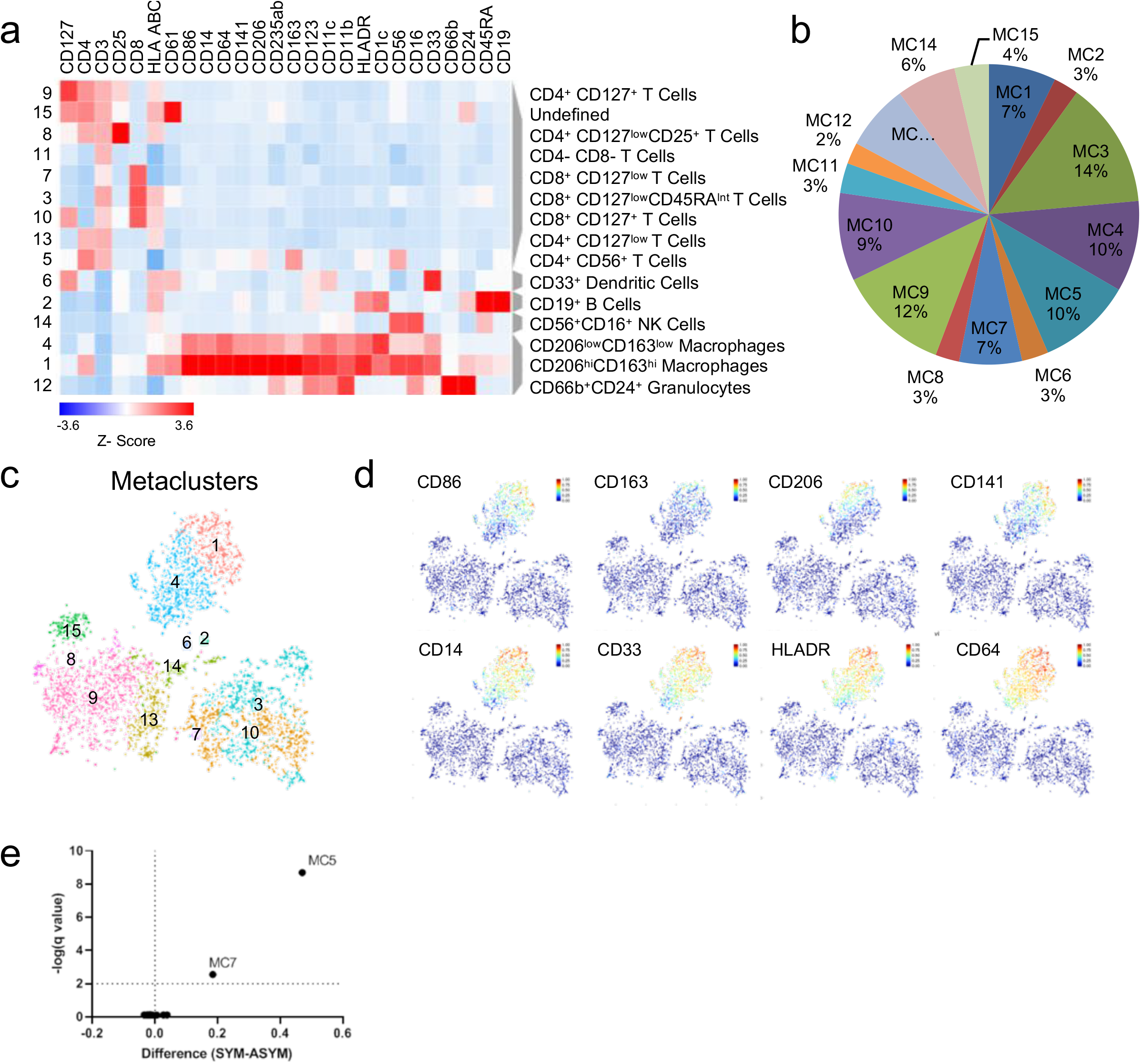
Tissue–specific Metaclustering Identifies Dysregulations in T Cells. (a) MetaLouvain Clustering of CD45^+^ cells from atherosclerotic plaque tissue (n=15). (b) Average population frequency of plaque-derived MCs. (c) Representative viSNE plot depicting individual MC distribution. (d) viSNE plots of selected myeloid MC surface markers. (e) Volcano plot of the fold change of MC frequencies in ASYM (left) and SYM (right) atherosclerotic plaques. Statistical significance was determined using unpaired t-test with multiple comparisons corrected by FDR=0.5%.

**Extended Figure 3.**
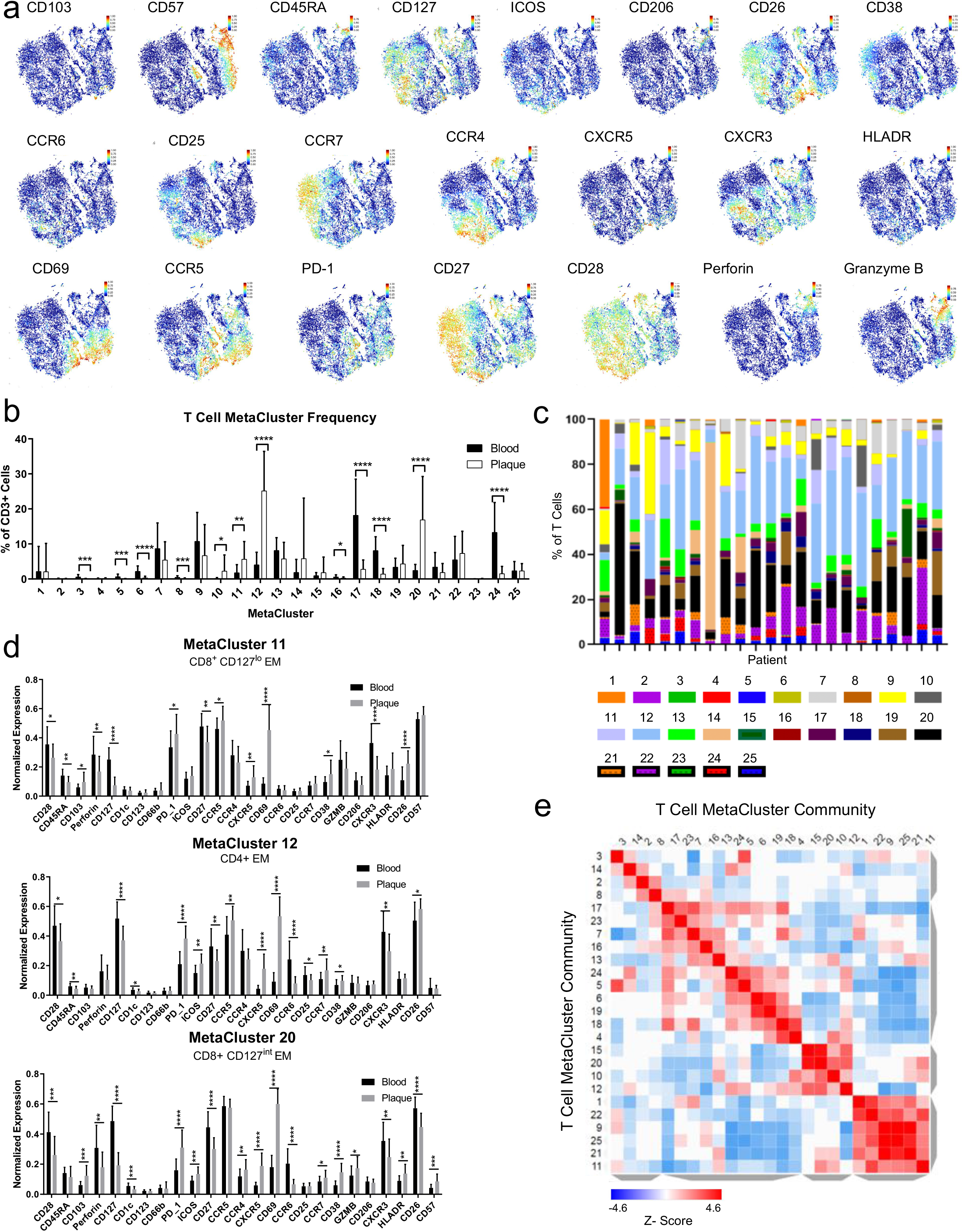
Definition of the T Cell Compartment from the Validation Cohort (cohort 2). (a) viSNE plots of T-cell MC data displaying the protein marker overlaid in spectral colors. (b) Bar plot of T-cell MC frequencies in blood and plaque. (c) Bar plot of MC frequencies per patient (n=23). (d) Bar charts of normalized surface marker expression of plaque-enriched MCs in blood and plaque. (e) Similarity matrix of T cell MetaClusters based on the cosine distance method. Statistical significance for (b) and (d) were determined using multiple unpaired Student’s t-tests corrected by the FDR=0.5%. *q<0.05, **q<0.01, ***q<0.001, and ****q<0.0001.

**Extended Figure 4.**
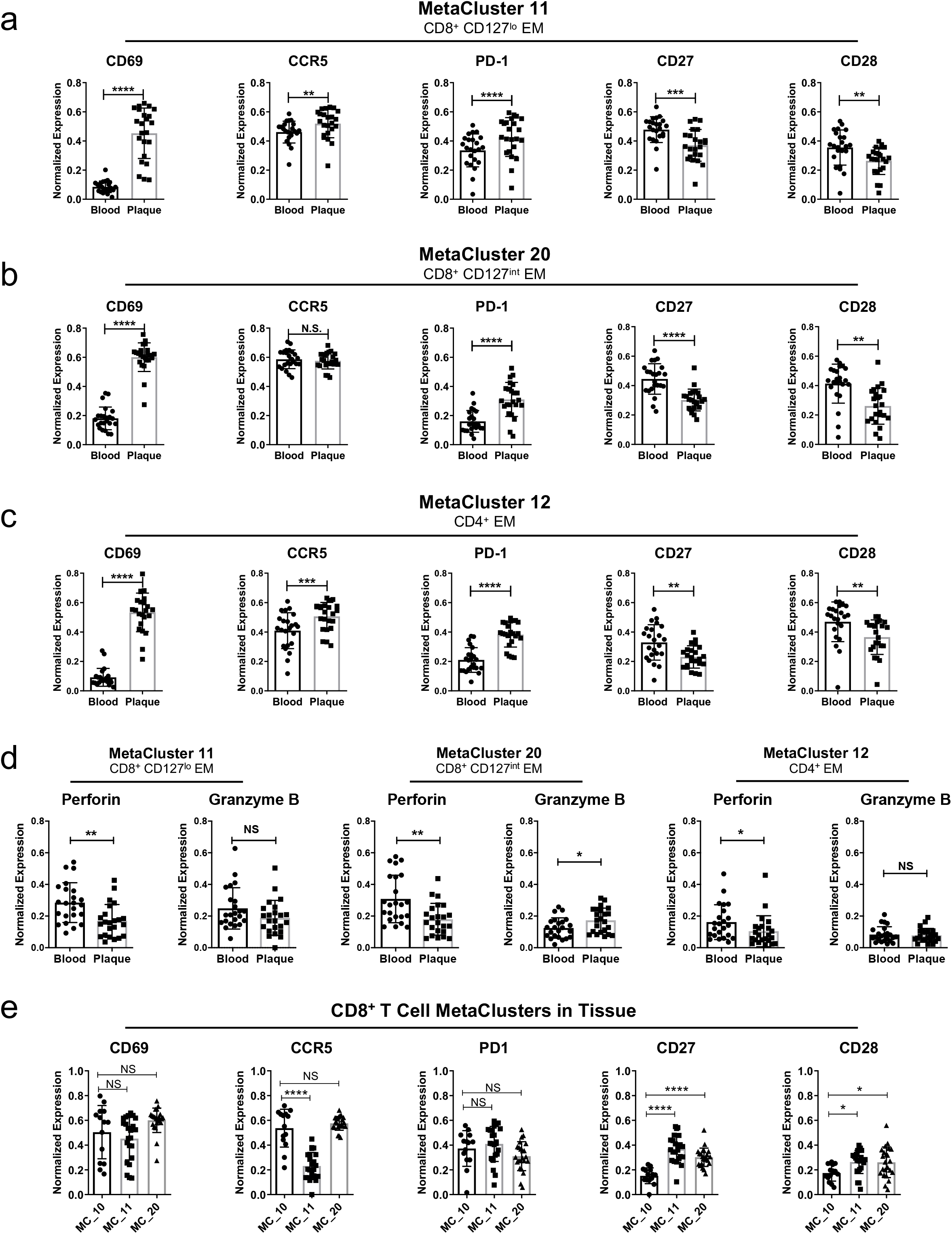
Unique T-Cell Signature in Atherosclerotic Plaques. Scatter bar plots of normalized marker expression between blood and plaque in the CD8^+^ and CD4^+^ T-cell MCs enriched in plaque (a-d), and comparison of MCs 10, 11, and 20 in plaque tissue (e). *p<0.05, **p<0.01, ***p<0.001, ****p<0.0001 by paired two tailed Student’s t-test (a-d), and One-Way ANOVA test with Bonferroni’s post-hoc correction (e). Values are mean ± SD.

**Extended Figure 5.**
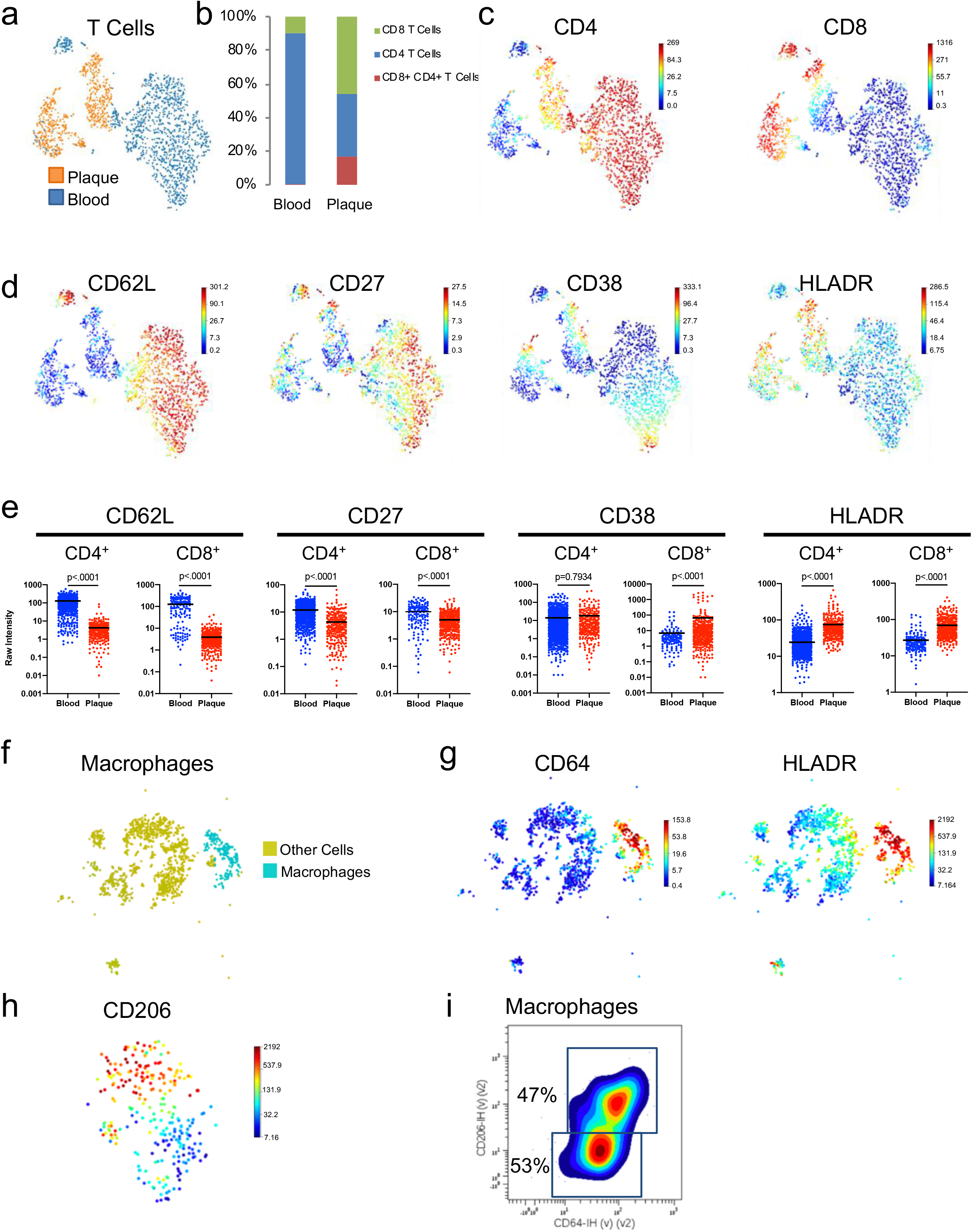
Analysis of the CITE-seq ADT Expression. viSNE plots of T cells clustered using the ADT data, and overlaid by (a) tissue type, or expression of canonical (c) or functional (d) T cell markers. (b) Frequency of T cell subtypes per tissue. (e) Functional surface marker expression in individual cells per tissue type. P values were determined using the Mann-Whitney test, lines indicate average expression values. (f, g) viSNE plots of plaque cells overlaid to indicate macrophage population (f), or expression of macrophage markers (g). (h) viSNE plot of clustered macrophages overlaid with CD206 expression. (i) macrophage population gated for expression of CD206 and CD64.

**Extended Figure 6.**
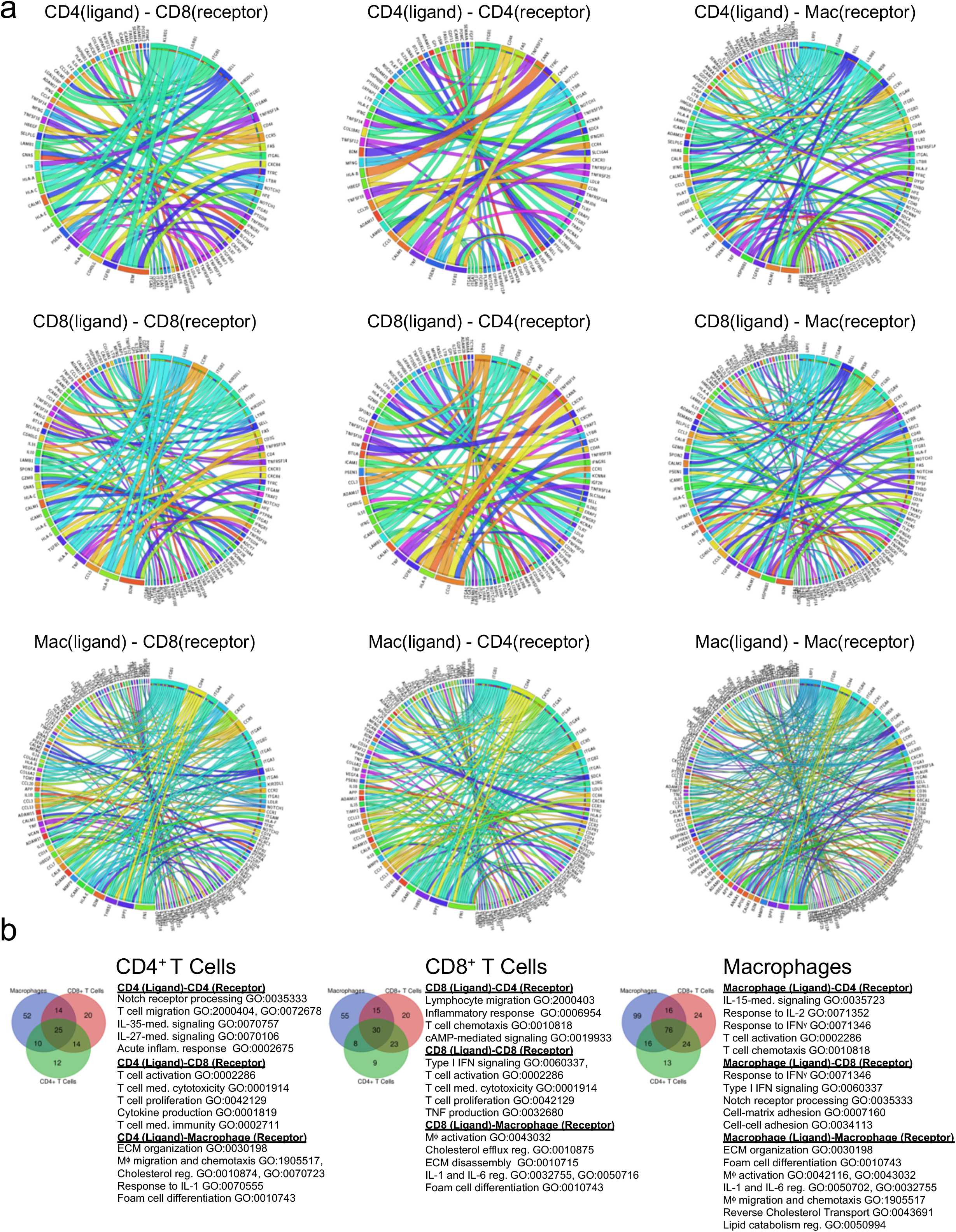
All Protein-Ligand Interactions. (a) Circos plots of the significant ligand-receptor interactions between cell types, mediated by CD4^+^ T cells (top row), CD8^+^ T Cells (middle row), or macrophages (bottom row). (b) Venn diagrams of ligand-receptor pairs from the top 5000 genes (>0.5 Log2 fold change) show unique and overlapping paired between cell-types. Gene Ontology terms were identified for each group using *Enrichr*.

## METHODS

### Human Specimens and Patient Clinical Characteristics

Forty-six patients undergoing carotid endarterectomy at the Mount Sinai Hospital were enrolled in an ongoing clinical study approved by the Institutional Review Board of the Icahn School of Medicine at Mount Sinai (IRB 11-01427). Eligible subjects gave informed written consent. The exclusion criteria were current infection, autoimmune disease, active or recurrent cancer, severe renal failure requiring dialysis, peripheral arterial occlusive disease causing pain at rest. Peripheral venous blood and atherosclerotic specimens were obtained from each patient.

Supplementary Table 1 summarizes the demographic and clinical characteristics of the patients. There were 29 men and 17 women with a mean age of 72.9±8.7 years (median 73.0 years). Symptomatic patients were defined as having had a cerebral ischemic event (stroke, TIA, amaurosis fugax) ipsilateral to the collected plaque within 6 months according to validated criteria^157,158^. Asymptomatic patients had no ischemic events within 6 months before surgery.

### Sample Collection

Fasting blood was collected preoperatively into tubes containing ACD Solution A as anti-coagulant (BD Vacutainer, 364606). Carotid atherosclerotic tissue specimens were obtained at the bifurcation point and placed immediately in Dulbecco’s modified Eagle’s medium (DMEM, Corning, 10-013-CV) supplemented with 10% fetal bovine serum (FBS, Gibco, 10082-147) on ice.

### Isolation of PBMCs

Blood was processed within 2 hr of collection and spun at 1500 rpm for 10 min using an Eppendorf A-4-62 rotor. Plasma was collected, and the cell fraction was diluted 1:2 with phosphate-buffered saline (PBS, Corning, 1-031-CV), placed on Ficoll-Paque Plus (ratio 7:3) (GE Healthcare Life Sciences, 17144003), and centrifuged for 20 min at 1800 rpm at room temperature. After centrifugation, the PBMC layer was collected and washed twice in PBS at room temperature.

### Cell Dissociation from Atherosclerotic Plaques

Plaque specimens were processed within 1 hr after surgery. Each specimen was washed extensively in DMEM, weighed, and digested as described 168 at 37° C for 1 hr in 10 ml of DMEM containing 10% fetal bovine serum; collagenase type IV (Sigma, C5138) at a final concentration of 1 mg/ml; and DNase (Sigma, DN25), hyaluronidase (Sigma, H3506), collagenase type XI (Sigma, C7657), and collagenase type II (Sigma, C6885), each at a final concentration of 0.3 mg/ml. The mixture was filtered consecutively through 70 μm and 40 μm cell strainers, washed twice in PBS, and centrifuged at 300 g for 8 min. Dead cells were removed with a kit (Miltenyi Biotec, 130-090-101, or Stem Cell Technologies, 17899) according to the manufacturer’s instructions. Cells were counted with an automatic cell counter.

### Preparation of Paired Samples for CyTOF

Cells isolated from plaque specimens and paired PBMCs from the same patient were barcoded using a CD45-based approach and optimized for scarce clinical samples^159^, pooled, and stained with antibody panels. All antibodies were validated at the Human Immune Monitoring Center of the Icahn School of Medicine at Mount Sinai. Antibodies were either purchased pre-conjugated from Fluidigm, or purchased purified and conjugated in-house using MaxPar X8 Polymer Kits (Fluidigm) according to the manufacturer’s instructions. The samples were then washed and stained with cisplatin-195Pt or Intercalator Rh103 (Fluidigm, 201064 and 201103A) as a viability dye^160^, washed, fixed, and permeabilized with BD Perm/Wash and Cytofix/Cytoperm buffers (BD Biosciences, 51-2090KZ and 51-2091KZ), and stored in freshly diluted 2% formaldehyde (Electron Microscopy Sciences) in PBS containing 0.125 nM Iridium 191/193 intercalator (Fluidigm, 201192) until acquisition. See Supplementary Table 3 for list of antibodies used in this study. A limitation of our T cell focused panel consisted in the lack of anti-foxp-3 antibody, which limited the ability of dissecting the Tregs compartment in human atherosclerosis.

### Acquisition and Processing of CyTOF Data

CyTOF data were acquired with a CyTOF2 using a SuperSampler fluidics system (Victorian Airships) at an event rate of <400 events per second and normalized with Helios normalizer software (Fluidigm). Barcoded samples were deconvoluted and cross-sample doublets were filtered using a Matlab-based debarcoding application^44^. Data were uploaded to Cytobank (https://mtsinai.cytobank.org; Cytobank, Menlo Park, CA) for analysis and visualization.

### Preprocessing of CYTOF Data

CYTOF data were processed with Cytobank to sequentially remove calibration beads, dead cells, debris, and barcodes for CD45+ plaque and CD45+ blood cells. For Cohort 1, we analyzed 41,222 plaque cells with an average of 2,748 cells per sample. For Cohort 2, we analyzed 49,088 plaque cells with an average of 2,134 cells per sample. For the T-cell MC analysis (Figure 3) and manual gating analysis (Supplementary Data to Figure 3), T cells were considered as CD3^+^CD14^lo^CD16^lo^CD56^lo^CD19^lo^CD64^lo^, and gated as such. For the manual gating analysis, T cells were further stratified as CD8^+^ or CD4^+^. Sub-classifications were defined as follows: naïve (CD45RA^+^CCR7^+^) CM (CD45RA^−^ CCR7^+^), EM (CD45RA^−^CCR7^−^), EMRA (CD45RA^+^CCR7^−^), T regulatory cells (CD4^+^CD127^lo^CD25^+^CCR4^+^), Th1 (CD4^+^CXCR3^hi^CCR6^−^), Th2 (CD4^+^CXCR3^lo^CCR6^+^), and Th17 (CD4^+^CXCR3^lo^CCR6^hi^) (Supplementary Table 4)^53^.

### Visualizing and Clustering CyTOF Data

CyTOF data were analyzed and visualized with Cytobank. Tissue-associated inflammatory cells and matched PBMCs were analyzed with an R-based semi-automated pipeline developed at the Human Immune Monitoring Center of the Icahn School of Medicine at Mount Sinai. This pipeline uses viSNE^43^ as a dimensionality-reducing visualization tool and Phenograph^44^ as an automated clustering algorithm. Stratifying signatures between the sample groups were identified by calculating the fold change between mean cluster abundances across groups. P values were calculated using a paired, two-sided Student’s t-Test using the FDR correction of the R package. Fold change and p values were visualized using a volcano plot (See Figures 2 and 3, and Supplementary Table 2). Clusters were manually annotated according to canonical patterns of marker expression, and Marker expression values were Z-scored and visualized with Clustergrammer (http://amp.pharm.mssm.edu/clustergrammer/). Statistical analysis of cell population frequencies and marker expression using the multiple t test with FDR correction, one-Way ANOVA test with Bonferroni’s post-hoc correction, two tailed Student’s t-test, Wilcoxon test and Spearman rank correlation were done with GraphPad Prism 8.0.

### Characterization of Plaque Type

The carotid plaque specimens were fixed in formalin (Thermo Scientific, 9990244), embedded in paraffin at the Pathology Biorepository at Mount Sinai Hospital, cut into 5-μm-thick sections, placed on charged glass slides, and stained with Masson’s trichrome stain as recommended by the manufacture (American Master Tech KTMTRPT). The images were used to evaluate the plaque type according to the American Heart Association Classifications^84^.

### TCR Variable Beta Chain Sequencing and Statistical Analysis

The CDR3 variable regions of the TCRβ chain were sequenced with the ImmunoSEQ Assay (Adaptive Biotechnologies, Seattle, WA). Genomic DNA was extracted from formalin-fixed, paraffin-embedded tissue samples and amplified by bias-controlled multiplex PCR, followed by high-throughput sequencing. Sequences were collapsed and filtered to identify and quantitate the absolute abundance of each unique TCRβ CDR3 region^161^. Clonality was defined as 1 – Peilou’s eveness^162^ and calculated as described^41^. Clonality values range from 0 to 1 and describe the shape of the frequency distribution: clonality values approaching 0 indicate a very even distribution of frequencies, whereas values approaching 1 indicate an increasingly asymmetric distribution in which a few clones are present at high frequencies. Correlations between clonality and CyTOF data were assessed with Spearman’s rank correlation test after running a Shapiro-Wilk test for normality. Statistical analyses were done with GraphPad Prism 8.0.

### Cellular Indexing of Transcriptomes and Epitopes by sequencing (CITE-seq) and Single-Cell RNA sequencing (scRNA-seq) Sample Preparation

A protocol adapted from Stoeckius et al 2017^42^ was used to perform CITE-seq. All antibodies used for CITE-seq were conjugated at the Human Immune Monitoring Core at Mount Sinai using ThunderLink Plus conjugation kits (Expedeon, Cat#425-0000), following manufacturer’s instructions. Single cell suspensions of atherosclerotic plaque and PBMCs were enriched for immune cells using a CD45^+^ enrichment kit (Miltenyi, 130-045-801), and equal proportions of CD45^+^ enriched blood and plaque samples were barcoded and immuno-stained on ice with the CITE-seq antibody panel (See Supplemental Table 3).

The single-cell suspensions of CITE-seq and scRNA-seq samples were converted to barcoded scRNA-seq libraries by using the Chromium Single Cell 3’ Library, Gel Bead & Multiplex Kit, and Chip Kit (10x Genomics)^163,164^. Chromium Single Cell 3’ v2 Reagent (10X Genomics, 120237) kit was used to prepare single-cell RNA libraries according to the manufacturer’s instructions. DNA quality was measured with an Agilent Bioanalyzer and quantity was assessed with the Qubit fluorometric assay (Invitrogen). Both CITE-seq and scRNA-seq used Cell Ranger Single-Cell Software Suite (version 2.1.1, 10X Genomics) to quality control the single-cell expression data. For CITE-seq, the Cell Ranger outputs estimated 3,573 cells with 50,701 mean reads per cell, and 3,879 median UMI counts per cell. For the scRNA-seq data a total of 7,169 cells were analyzed and averages of the Cell Ranger outputs from the n=6 patients an average of 111,670 mean reads per cell, and 2,929 median UMI counts per cell. The filtered gene-barcode matrix output from Cell Ranger, which removes barcodes with a low unique molecular identifier (UMI) count, was analyzed in further analyses using Clustergrammer2 (https://github.com/ismms-himc/clustergrammer2)^165^.

### CITE-seq Expression Analysis

PBMC and Plaque-derived cells were identified by de-hash tagging (following the cell hashing technique from Stoeckius et al^42^). The surface marker expression, from Antibody derived tags (ADT) data from a 21 antibody panel (see Supplementary table 3) was analyzed using Cytobank. First, doublets between PBMCs and plaque were removed by exclusion of overlapping barcodes. Second, all individual cells were clustered using the t-SNE algorithm based on expression of all 21 markers. Third, immune cell types were determined based on the expression of canonical markers, and were thus gated directly from the viSNE plot. Correlations of the gene and surface marker expression can be found in Supplementary Figure 9.

Ribosomal and mitochondrial genes were dropped from the gene expression analysis. Gene expression signatures of the atherosclerotic plaque and blood were derived from the cell types obtained from the ADT manual gating (CD4 T cells, CD8 T cells, B Cells, Macrophages, Monocytes, NK Cells, NKT Cells, Plasma Cells, CD1c DCs, pDCs), and used to predict cell type across the cohort of single-cell RNA-seq of atherosclerotic plaques. The gene expression data corresponding to the cells of gated populations was used for further analysis by *Clustergrammer2*, *Enrichr*, and Ingenuity Pathway Analysis (IPA).

### Batch Correction of Single-Cell RNA-seq data

Plaque gene expression from six subjects was combined, and batch normalized using mutual nearest neighbors (MNN) (https://github.com/chriscainx/mnnpy)^166^. First, gene expression data from all subjects was combined into a single dataset and suspected red blood cells were dropped (e.g. cells with high hemoglobin expression). Second, gene expression data was Arcsinh normalized and filtered to keep the top 10,000 genes based on variance across the aggregated dataset as described^166^. Third, MNN batch correction was performed using the one of the samples as a reference (See **Supplementary Figure 10**).

### Cell Type Prediction of Single-Cell RNA-seq using CITE-seq Gene Expression Signatures

Cell type from single-cell gene expression data across six subjects was predicted using a gene expression signature method. First, 1,641 single cells from the CITE-seq sample were assigned a cell type based manual annotation of single cell t-SNE clustering in surface marker space (cell types include: CD4^+^ T cells, CD8^+^ T cells, B Cells, Macrophages, Monocytes, NK Cells, NKT Cells, Plasma Cells, CD1c DCs, pDCs). Second, cell type gene expression signatures were generated from the MNN batch corrected CITE-seq gene expression data. Cell type signatures were calculated by Z-scoring gene expression levels across all 7,169 cells, averaging gene expression levels across cell types, and keeping the top 200 differentially expressed genes for each cell type (using the Welch’s T-test). Third, cell type is predicted for all 7,169 cells across all samples by assigning each cell to the closest cell type signature based on cosine distance (gene expression data is Z-scored before distance calculation). Cell type prediction of the CITE-seq dataset using the MNN-batch corrected gene expression signatures matched previous ADT-based assignments 81% of the time.

### Single-Cell RNA-seq and CITE-seq Gene Expression and Pathway analysis

Single-cell gene expression data was analyzed for differential genes expression across tissue types (CITE-seq), or across patient types (using aggregated scRNA-seq). Differential gene expression was calculated using UMI normalized gene expression data.

When differential expression was calculated using data from multiple subject samples, which had been subject to MNN batch correction, UMI normalized gene expression data was used in place of MNN normalized data as described^166^. Differential expression was calculated using the Welch’s T-test with multiple hypothesis correction using Benjamini-Hochberg correction. Single-cells were hierarchically clustered using the top genes based on variance to identify subclusters within specific cell types. The hierarchical clustering was done using cosine distance with average linkage. Clusters were defined by assigning groups at a manually chosen level in the dendrogram. Differentially expressed genes across cell clusters were identified with the Welch t test with unequal variance; multiple hypothesis correction was done with the Benjamini-Hochberg correction in the SciPy Stats package, (https://docs.scipy.org/doc/scipy/reference/index.html). For our canonical pathway analysis, we analyzed cell clusters containing at least 25 cells using Ingenuity Pathway Analysis (QIAGEN Inc.). The top 5000 DEGs corresponding to the identified cell clusters were investigated with the gene enrichment analysis tool *Enrichr*^167,168^. For CITE-seq, the Enrichr pathway analysis terms were chosen from the Reactome library. Reactome, Biocarta, NCI Nature, and GO Biological Functions libraries were used to analyze the scRNA-seq data. Terms from these libraries were plotted using the combined score, which represents the p value (Fisher’s exact test) multiplied by the Z-score of the deviation from the expected rank.

### Cell Type Ligand-Receptor Interaction

Potential ligand-receptor interaction between one set of ligand expressing cells and another set of receptor expressing cells is calculated as the average of the product of ligand and receptor expression (respectively from set one and two) across all single-cell pairs:

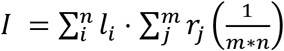

*I*= interaction score between ligand expressing cells in set one and receptor expressing cells in set two
*I_i_* = ligand expression of cell *i* in cell set one
*r_j_* = receptor expression of cell *j* in cell set two
*n*= number of cells in set one
*m*= number of cells in set two

Prior knowledge ligand and receptor interactions were obtained from Ramilowski et al^169^. Cell types were obtained from the 7,169 cell MNN batch corrected dataset and gene expression levels were calculated using UMI normalized data. Potential ligand-receptor interactions were calculated for three cell types (CD4 T Cells, CD8 T Cells, and Macrophages) resulting in nine cell-type/cell-type interaction scores. Visualization of the interactions was performed using CIRCOS plots (http://mkweb.bcgsc.ca/tableviewer/), *and Clustergrammer*. Comparison of receptor-ligand pairs between cell types were analyzed visualize using a Venn diagram generate from (http://bioinformatics.psb.ugent.be/webtools/Venn/).

Differential regulation of ligand-receptor interactions between symptomatic and asymptomatic cells was identified by comparing single-cell/single-cell ligand-receptor interaction scores for symptomatic and asymptomatic cells. Fold change of symptomatic vs asymptomatic interaction scores were calculated using the average interaction scores from symptomatic and asymptomatic. The significance of differential regulation was assessed by comparing the distributions of cell-cell ligand-receptor interaction scores from symptomatic and asymptomatic cells using the Welch’s T-test. Symptomatic vs Asymptomatic differentially regulated interactions were identified based on the following 4 criteria: (1) one percent of cells must express the ligand and receptor of interest in either symptomatic or asymptomatic cells, (2) the interaction score of symptomatic or asymptomatic cells must be at least half of the baseline interaction score (defined as the interaction score between all cells in the dataset), (3) the log2 fold change of the interaction score (symptomatic vs asymptomatic) must be greater than 0.5, (4) the P-value of the Welch’s T-test with multiple hypothesis correction (Benjamini-Hochberg) must be less than 0.05. For visualization, log2 Fold Change values were clipped at +/-10.

### Data and Software Availability

The data discussed in this publication will be deposited using Kaggle Datasets and be publicly available.

