## Supplementary Figures for "Immune Profiling of Atherosclerotic Plaques Identifies Innate and Adaptive Dysregulations Associated with Ischemic Cerebrovascular Events"

### Supplementary Figure 1

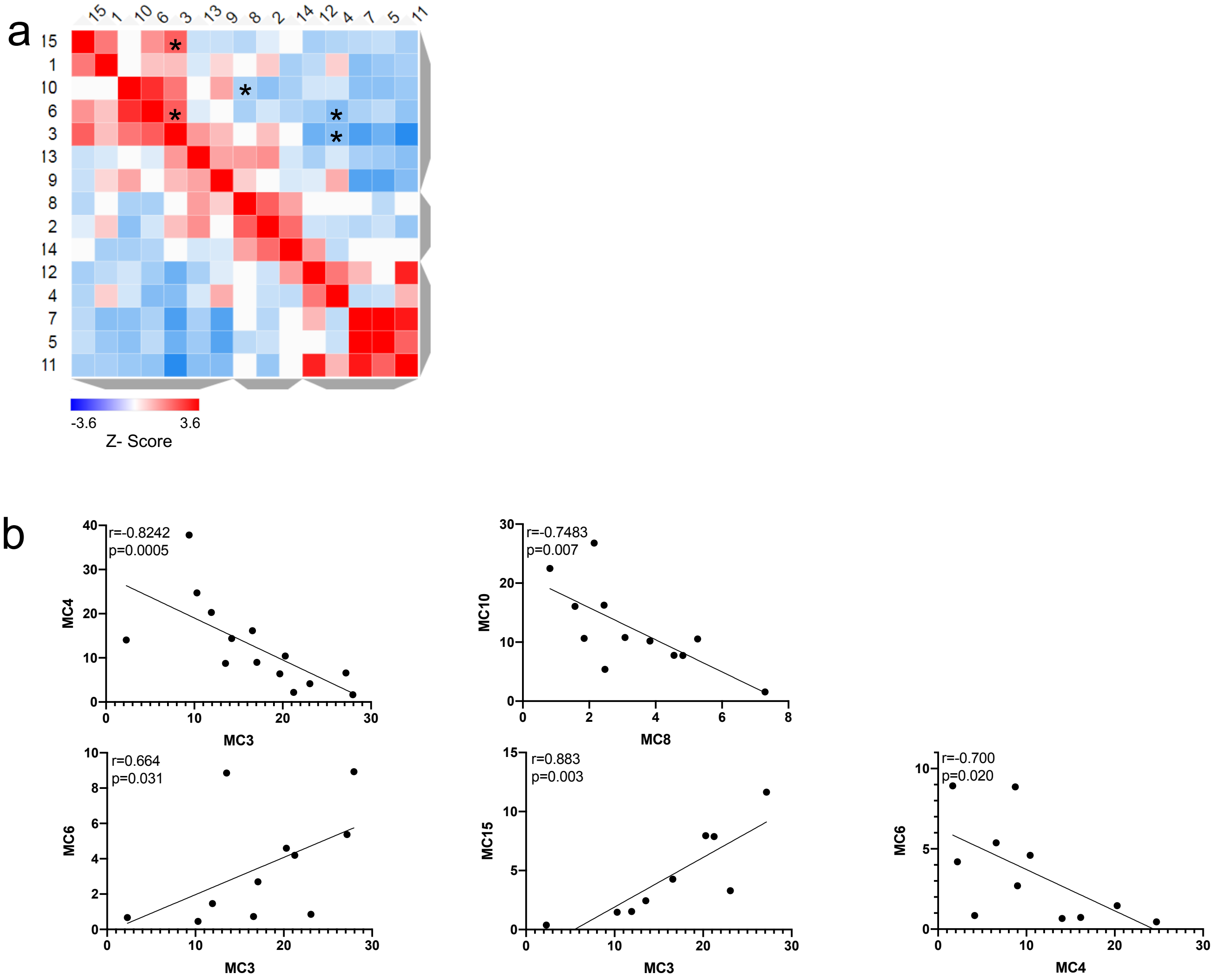

### Supplementary Figure 2

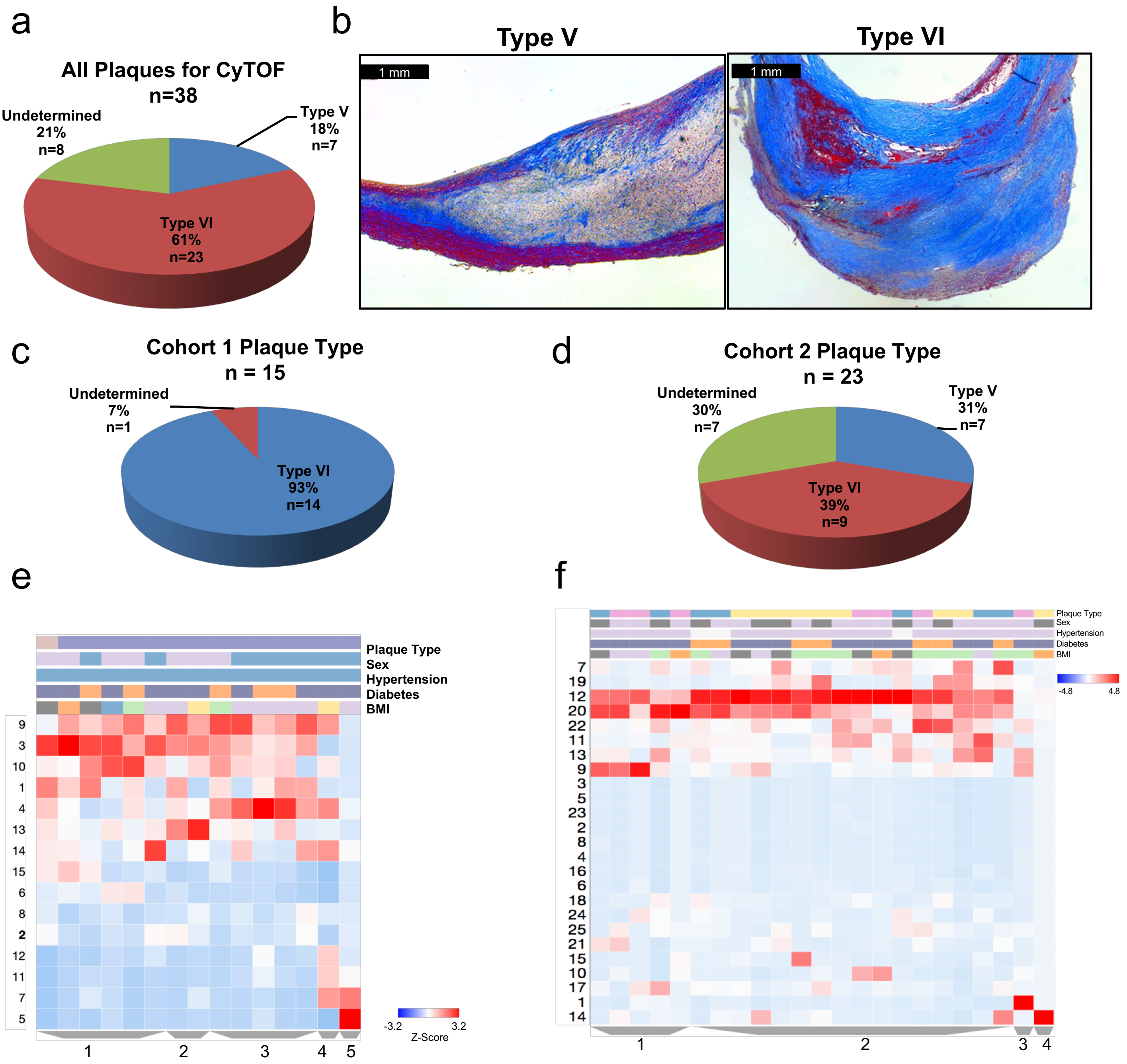


g

| Cluster | 1 | 2 | 3 | 4 | 5 |
|---|---|---|---|---|---|
| Plaque: Type VI | 0.94 | 0.87 | 0.71 | 0.93 | 0.93 |
| Undetermined | 34 | -------- | -------- | -------- | -------- |
| Sex: Male | 0.92 | -------- | 0.23 | 0.53 | 0.53 |
| Female | 0.28 | 0.22 | 0.96 | -------- | -------- |
| Hypertensive | 1 | 1 | 1 | 1 | 1 |
| Diabetic: Yes | 0.68 | 0.44 | 0.95 | 0.67 | 0.67 |
| No | 0.65 | -------- | 0.21 | -------- | -------- |
| BMI: Normal | 0.58 | -------- | 0.51 | -------- | -------- |
| Overweight | 0.98 | 0.72 | 0.15 | -------- | 0.47 |
| Obese | 0.19 | ------- | -------- | -------- | -------- |
| Extremely Obese | 0.34 | ------- | -------- | -------- | -------- |
| Underweight | -------- | ------- | -------- | -------- | -------- |
| Unknown | 0.34 | 0.25 | -------- | 0.13 | -------- |

h

| Cluster | 1 | 2 | 3 | 4 |
|---|---|---|---|---|
| Plaque: Type V | 0.17 | 0.57 | -------- | -------- |
| Type VI | -------- | 0.26 | -------- | 0.39 |
| Undetermined | 0.48 | 0.91 | 0.3 | -------- |
| Sex: Male | 0.7 | 0.56 | 0.61 | -------- |
| Female | 0.65 | 0.64 | -------- | 0.39 |
| Hypertensive: Yes | 0.5 | 0.85 | 0.87 | 0.87 |
| No | -------- | 0.35 | -------- | -------- |
| Diabetic: Yes | 0.16 | 0.92 | 0.7 | 0.7 |
| No | -------- | 0.19 | -------- | -------- |
| BMI: Normal | 0.94 | 0.39 | 0.43 | -------- |
| Overweight | 0.3 | 0.71 | -------- | -------- |
| Obese | 0.71 | 0.47 | -------- | -------- |
| Extremely Obese | 0.5 | 0.89 | -------- | 0.13 |

### Supplementary Figure 3

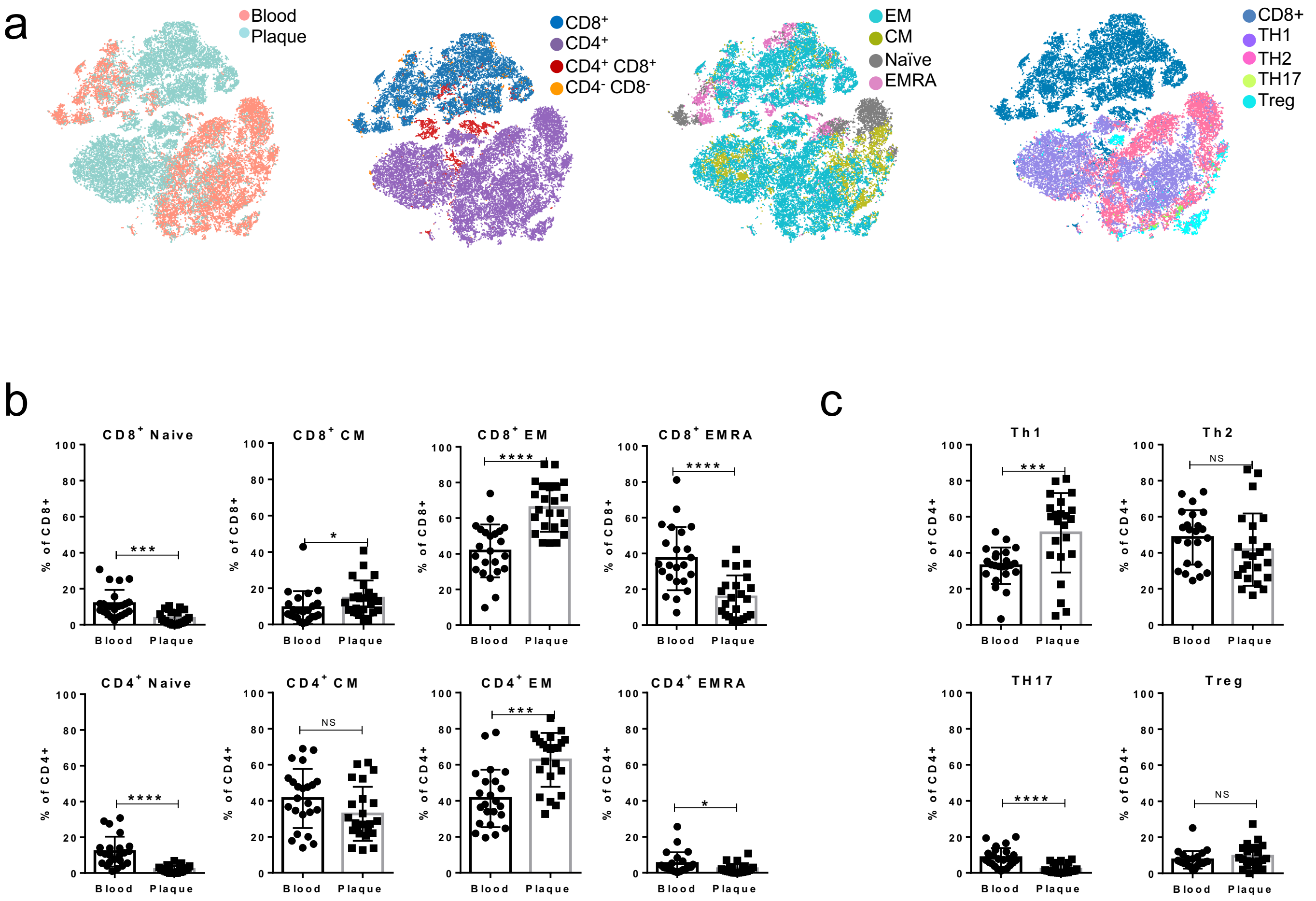

### Supplementary Figure 4

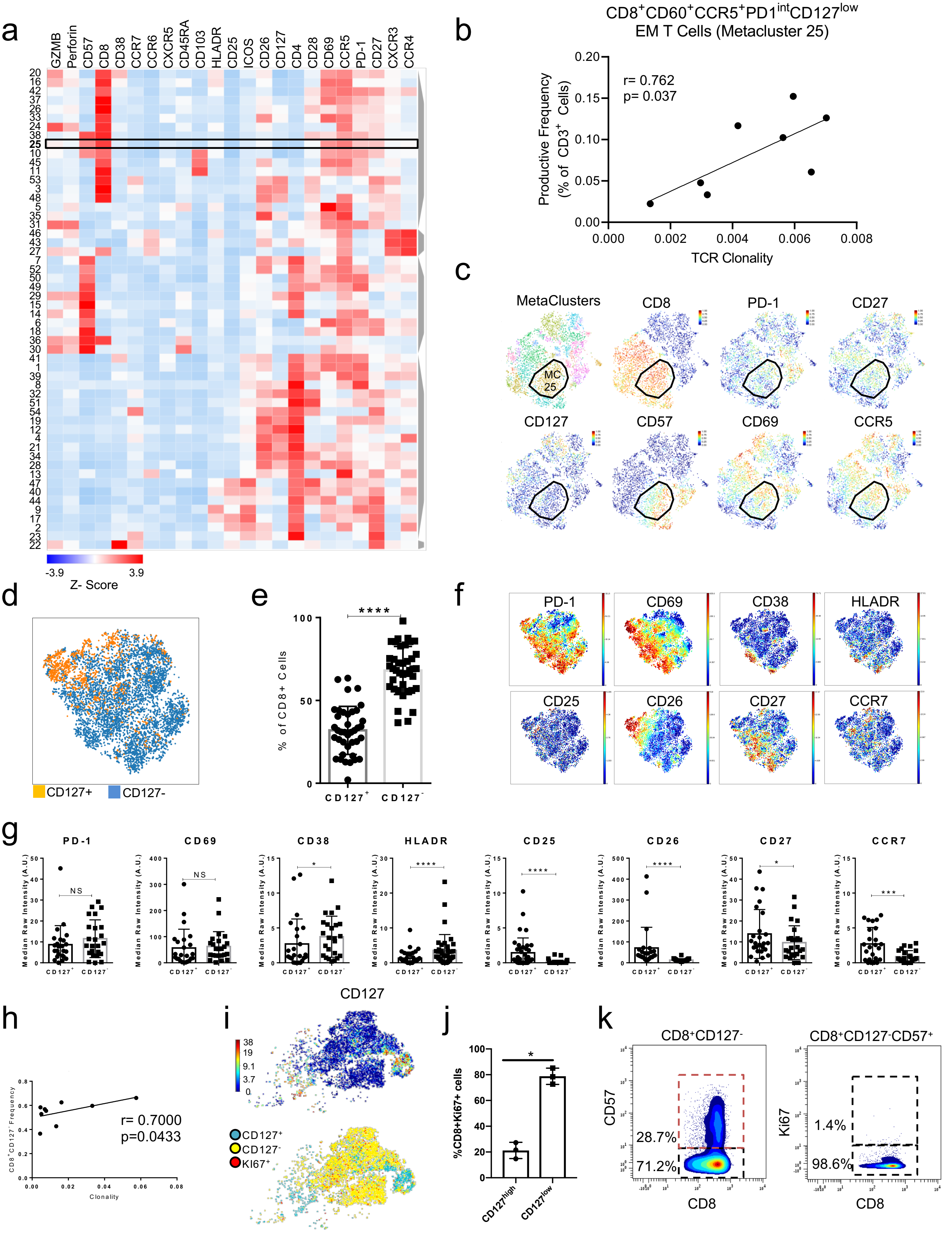

### Supplementary Figure 5

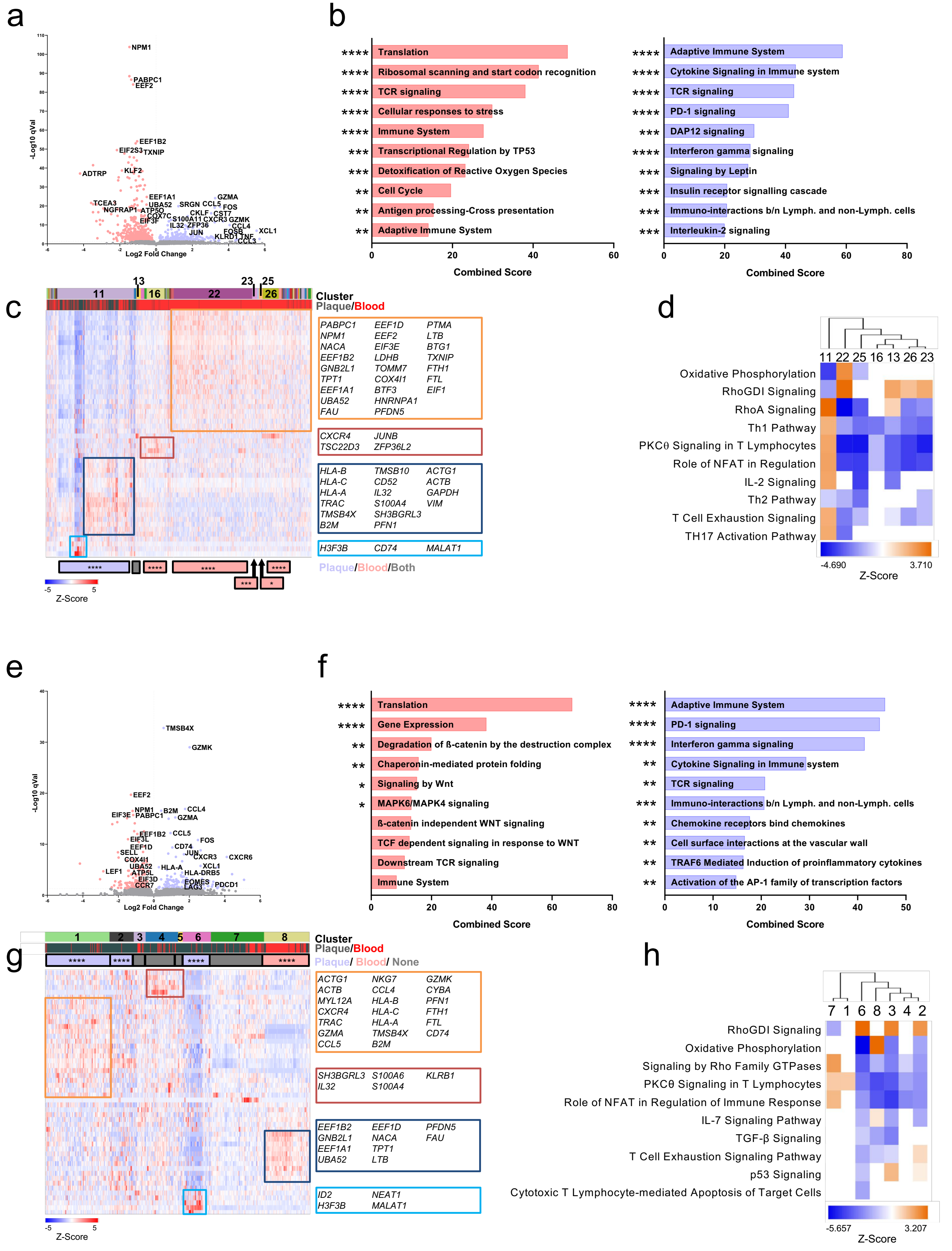

### Supplementary Figure 6

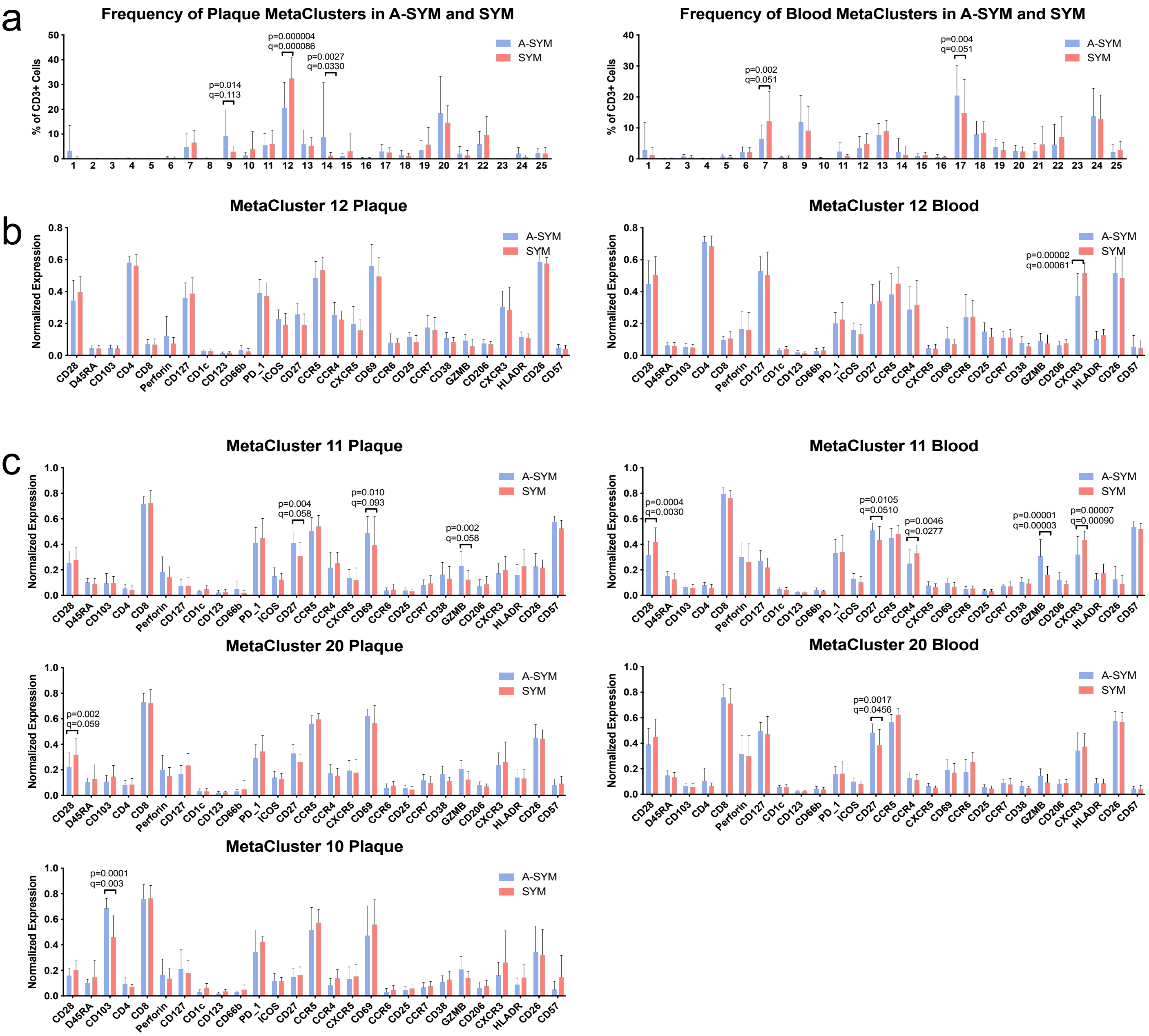

### Supplementary Figure 7

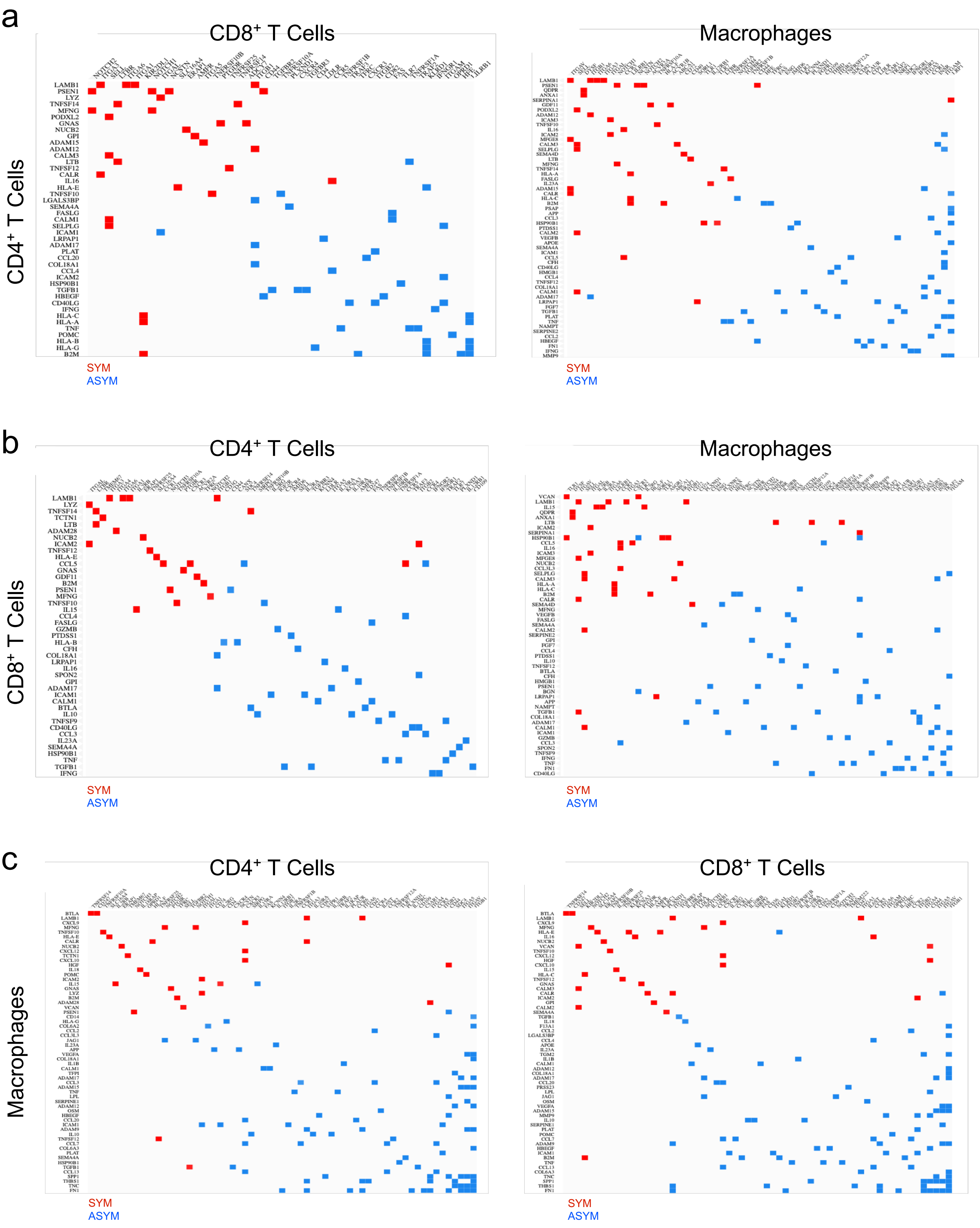

### Supplementary Figure 8

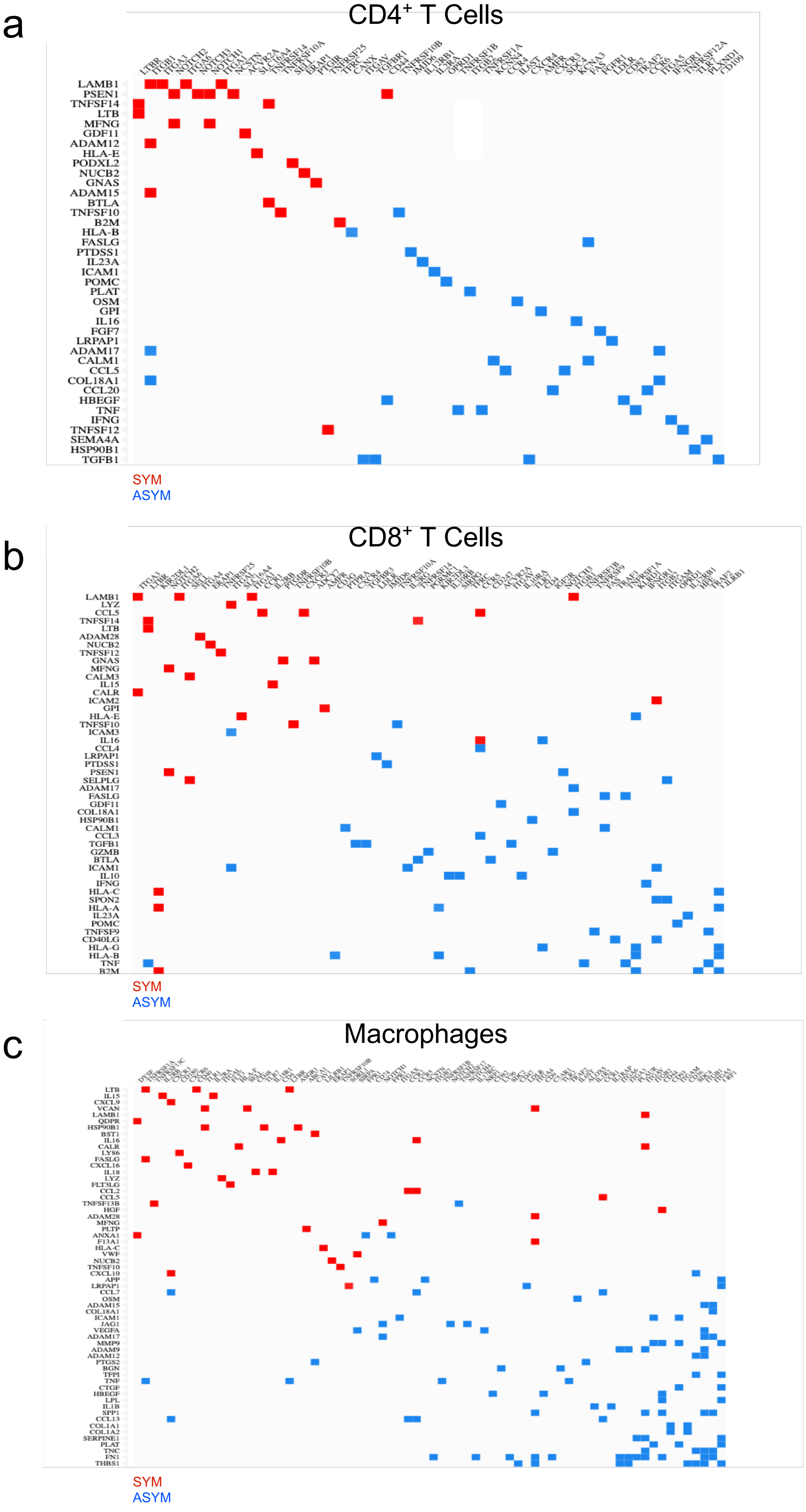

### Supplementary Figure 9

a

#### Blood

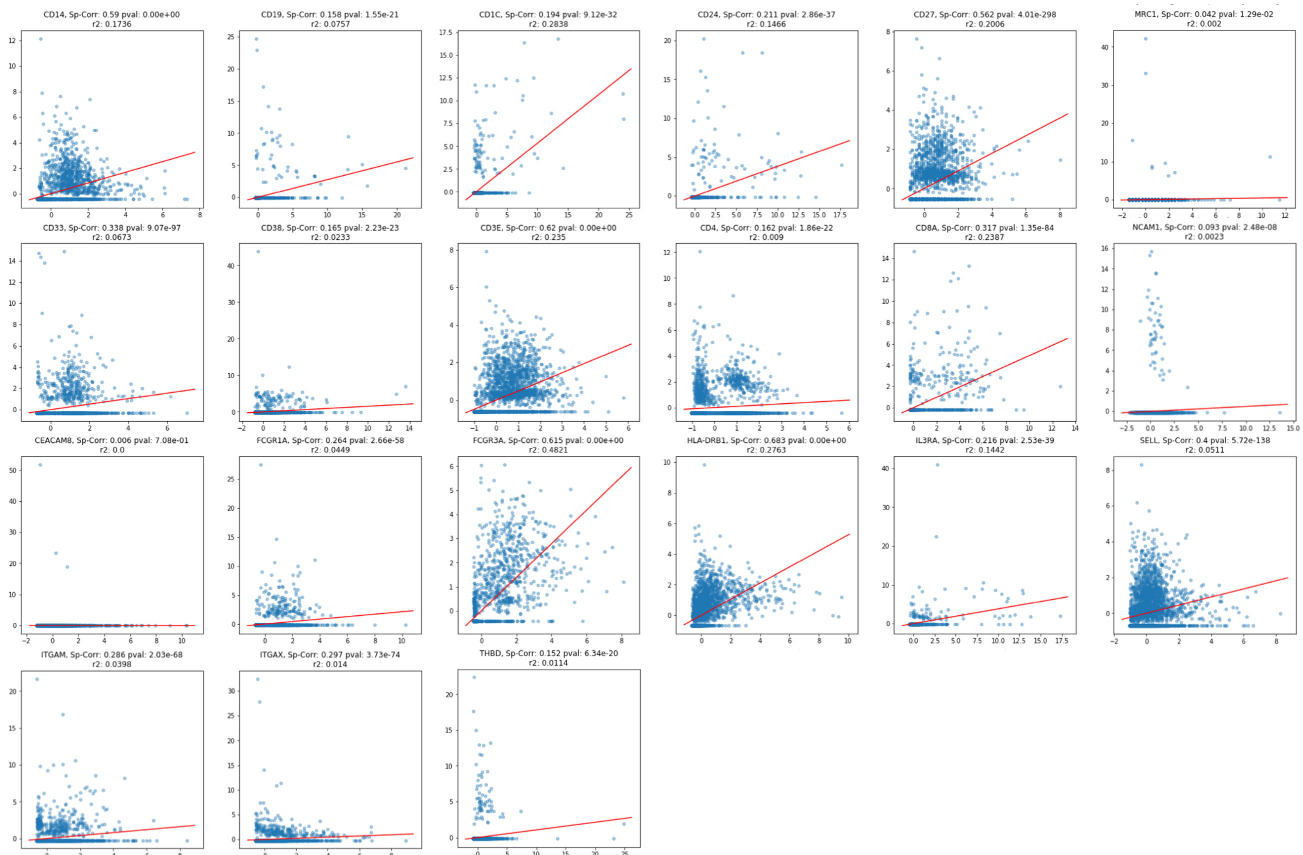


b

#### Plaque

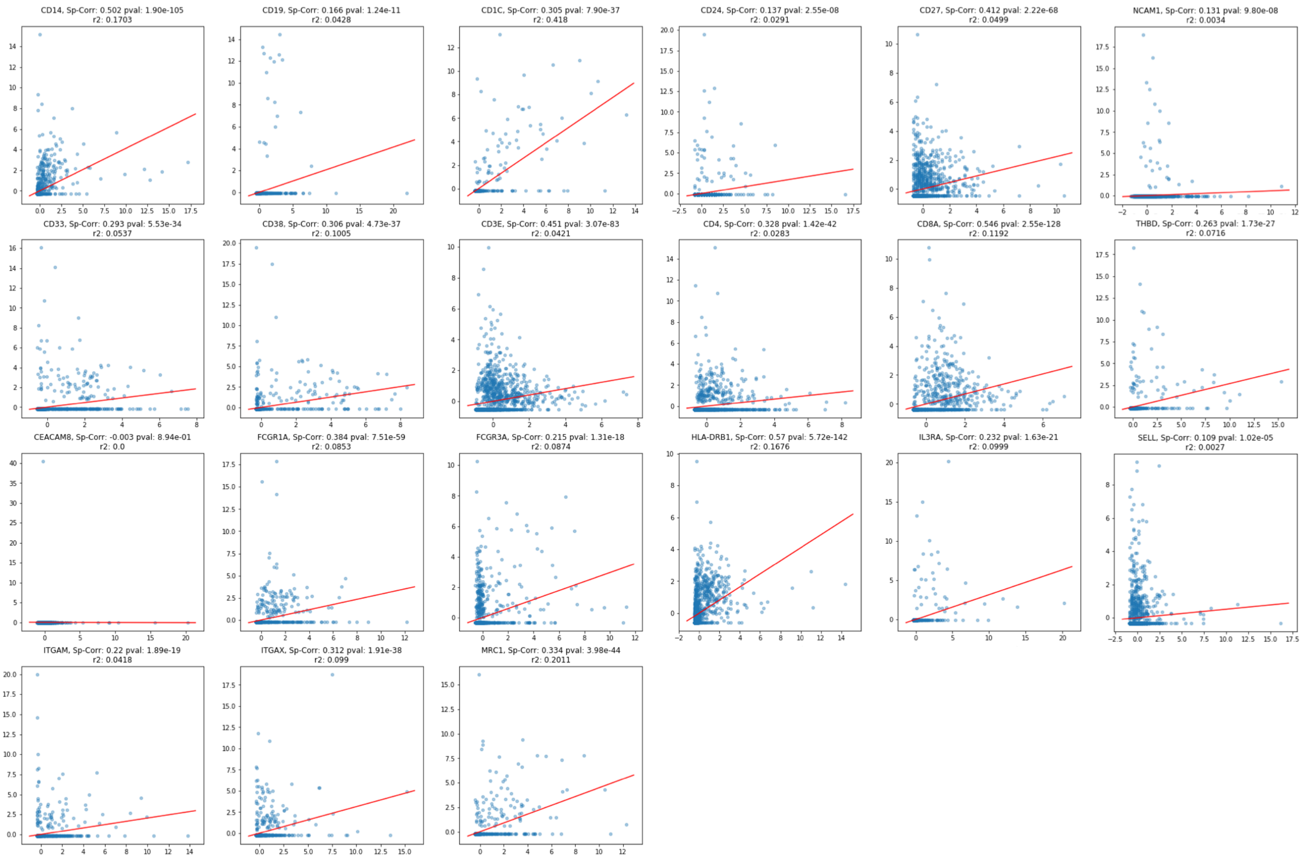

### Supplementary Figure 10

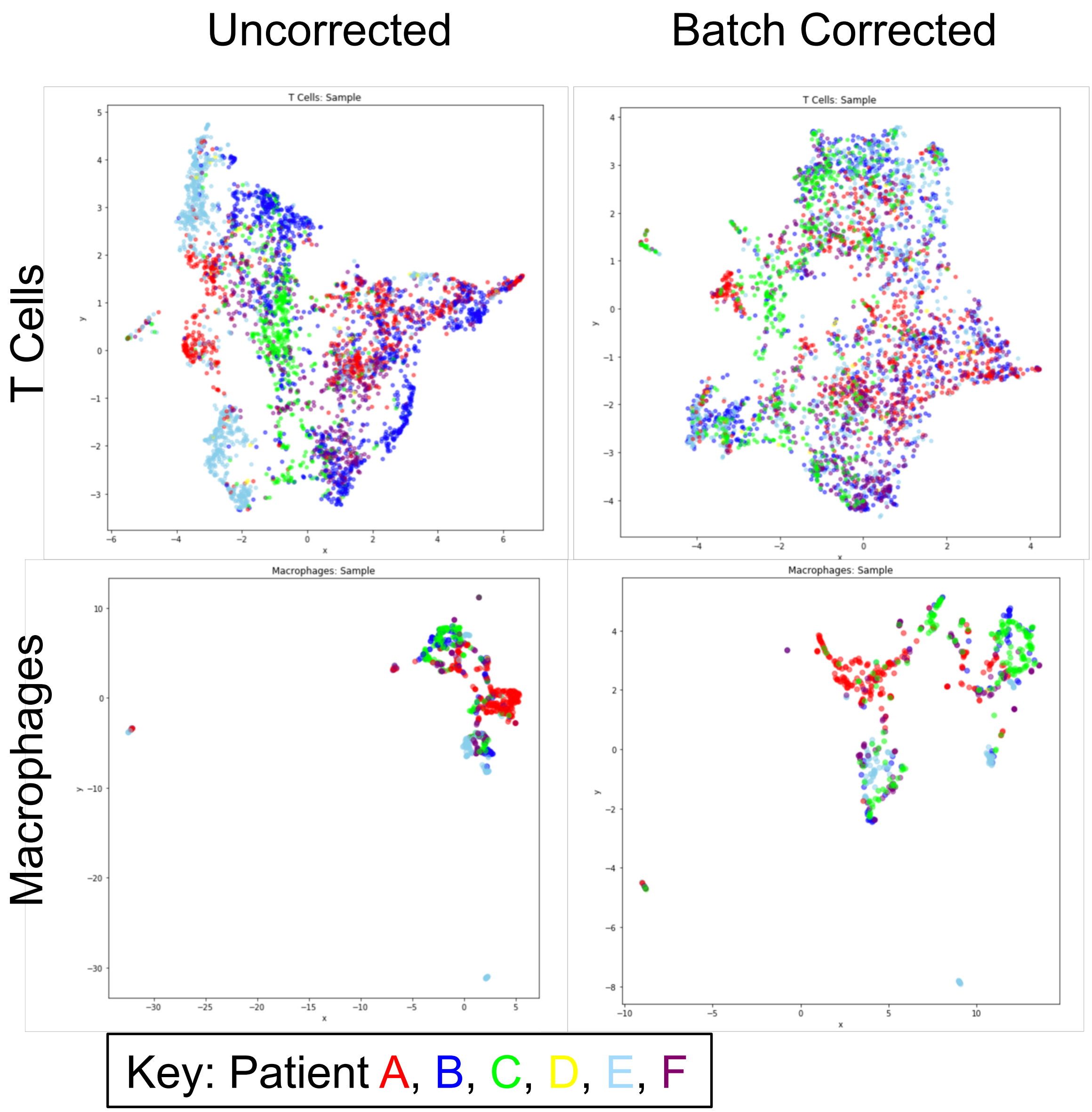
