## Supplementary Info for "Immune Profiling of Atherosclerotic Plaques Identifies Innate and Adaptive Dysregulations Associated with Ischemic Cerebrovascular Events"

### SUPPLEMENTARY INFORMATION

### SUPPLEMENTARY RESULTS

#### Single-Cell Mass-Cytometry of Immune Cells in Atherosclerotic Plaque.

We identified significant associations between cell frequencies of different plaque MCs suggesting dynamic interplay between the innate and adaptive immune composition in plaques. In particular, cell frequency of $CD8^{+}$ T cells of MC 3 was inversely associated with that of $CD163^{low}CD206^{low}$ macrophages of MC 4. Similarly, the frequencies of $CD8^{+}$ T cells of MC 10 and $CD4^{+}$ T cells of MC 8 were inversely associated, further suggesting that increases in $CD8^{+}$ T cell infiltrates are associated with lower macrophage and $CD4^{+}$ T cell subset content in plaques. Moreover, frequencies of $CD8^{+}$ T cells of MC 3 were positively correlated with dendritic cells of MC 6 and $CD4^{+}$ T cells of MC15. We also found an inverse correlation between $CD163^{low}CD206^{low}$ macrophages of MC 4 and dendritic cells of MC 6 (**Supplementary Fig. 1**).

#### Confirmation of Inflammatory T Cells Enriched in Plaque by Manual Gating

An independent manual gating analysis of T cells from n=24 patients confirmed that $CD8^{+}$ EM T cells were more abundant in atherosclerotic lesions than in blood and showed that most of the T cells in the lesions were $CD4^{+}$ and $CD8^{+}$ EM cells (**Supplementary Data Figure 3a, b)**. The gating strategy is reported in Supplementary Table 4. Additionally, since the $CD4^{+}$ T-cell compartment in atherosclerosis is traditionally defined as Th1, Th2, Th17, and regulatory T cells, we performed a second manual gating analysis (Supplementary Table 4). A limitation of this analysis consisted in the lack of an antibody against foxp3, a specific marker of Tregs, which limited the identification of Tregs to the the expression of CD25. We found that the majority of T cells in plaque were Th1 and Th2; however, only Th1 cells were more abundant in atherosclerotic lesions than in blood of the same patient (**Supplementary Data Figure 3a, c).**

#### Single-Cell Mass-Cytometry Analysis of T Cells Enriched in Atherosclerotic Plaques

Unbiased MC analysis of plaque T cells revealed two main categories of $CD8^{+}$ MCs based on the expression CD127, a marker of T cell differentiation, with 11 MCs of $CD127^{-}$ $CD8^{+}$ T cells compared to 3 subsets of $CD127^{+}$ $CD8^{+}$ T cells in plaques. Among all T cell MCs in plaque, we found a positive correlation between cell frequency of MC 25, a subset of $CD69^{+}CCR5^{+}PD1^{int}CD57^{int}CD27^{int}$ and $CD127^{-}$ $CD8^{+}$ T cells, and TCR clonality in tissue (**Supplementary Figure 4a-c)**. An independent manual gating analysis of plaque-specific $CD8^{+}$ T cells confirmed that the majority of $CD8^{+}$ subsets

(≈68%) were CD127$^{-}$ (**Supplementary Figure 4d-e**). Both CD127$^{-}$CD8$^{+}$ and CD127$^{+}$CD8$^{+}$ T cells expressed high levels of PD-1 and CD69; however, HLA-DR and CD38 expression were higher in the CD127$^{-}$ subset, indicating a higher activation state of these cells[1]. The CD127$^{-}$CD8$^{+}$ T cells also showed lower expression of CD26, CD27, and CCR7, suggesting a more differentiated phenotype[2] (**Supplementary Figure 4f, g**). As seen for the CD127$^{-}$ cells of MC25, we confirmed a positive correlation between CD127$^{-}$CD8$^{+}$ T cell frequency and TCR clonality in tissue (**Supplementary Figure 4h**), suggesting clonal expansion of these cells. Accordingly, CD127$^{-}$ T cells had a higher frequency of Ki67$^{+}$ cells than the CD127$^{+}$ subset (**Supplementary Figure 4i, j**), indicating a higher proliferative state of CD8$^{+}$CD127$^{-}$ T cells in atherosclerotic tissue. Of note, a subset of CD127$^{-}$CD8$^{+}$ T cells were CD57$^{hi}$ and ki67$^{-}$ (**Supplementary Figure 4k**), suggesting a senescent replicative transitioning possibly due to chronic antigen stimulation[3]. Thus, subsets of CD127$^{-}$CD8$^{+}$ EM T cells in atherosclerotic arterial wall appear to be clonally expanded, highly differentiated and transitioning to a senescent non-proliferative phenotype[3].

**Simultaneous Surface Marker and Gene Expression Immune Profiling Across Plaque and Blood: CITE-seq Analysis**

***GEX analysis of CD4$^{+}$ T cells in blood and atherosclerotic plaques***

We performed single-cell transcriptome analysis of n=1,830 CD4$^{+}$ T cells (**Supplementary Fig. 5a-d**). Consistent with our CyTOF data, circulating CD4$^{+}$ T cells were characterized by a resting phenotype while they presented an activated state in plaque. Genes upregulated in plaque were involved in Th1 functions (i.e. *KLRD1, KLRC1, CXCR3, STAT3, IFNGR1, HLA-B*) and chemotaxis (i.e. *CCL5, CCL4, CXCR6, XCL1*) and cytotoxicity (*GZMA, GZMK*), suggesting that subsets of CD4$^{+}$ T cells exhibit cytotoxic functions in plaques as in described in other disease states. Hierarchical clustering of CD4$^{+}$ T cells across plaque and blood identified a total of 26 clusters that were largely enriched in blood (**Supplementary Fig. 5c**). Among others, cluster 11, which was uniquely characterized by the expression of genes involved in T cell activation, T cell homing, inflammatory responses and was the only one associated with the specific upregulation of signaling pathways of T cell activation (i.e. *RhoA, PCKθ, NFAT*), differentiation and homeostasis of pro-inflammatory T cells [IL2 signaling, Th1 (*KLRD1, KLRC1, CXCR3, STAT3, IFNGR1, HLA-B0*), Th17 (*RORA, BATF*)], suggesting that these cells are highly inflammatory (**Supplementary Fig. 5d**). Interestingly, plaque cluster 11 also displayed the expression of genes associated with the pro-inflammatory Th17 signaling pathway and T cell exhaustion, indicating functional dysregulations of CD4$^{+}$ T cells as seen in other chronic inflammatory disease[4,5]. Clusters of CD4$^{+}$ T cells that were predominant in blood were characterized by a gene expression signature consistent with a resting phenotype (i.e. Oxidative

Phosphorylation and RhoGDI signaling), with the exception of Cluster 13 that also presented the upregulation of RhoA signaling thus indicating a certain degree of activation for these circulating cells (**Supplementary Fig. 5d**).

#### *GEX analysis of CD8$^+$ T cells in blood and atherosclerotic plaques*

A similar analysis (**Supplementary Fig. 5e-h**) of CD8$^+$ T cells confirmed that like for CD4$^+$ T cells, CD8$^+$ T cells were resting in blood and an activated in plaque. In blood, CD8$^+$ T cells were characterized by the upregulation of genes involved in β-catenin degradation, β-catenin-independent Wnt signaling (*PMSC5, LEF1, UBA52*) and downstream TCF signaling in response to Wnt (*TCF7, PSMD7, H3F3A*) (**Supplementary Fig. 5f**). An intact β-Catenin/Wnt/TCF signaling is implicated in the regulation of T cell development, but uncertain roles in the regulation of mature T cells, ranging from cell survival and pro-inflammatory functions to anergy, have been reported[6]. Our data indicate that this signaling is dysregulated in circulating CD8$^+$ T cells and suggest that these alterations may be implicated in determining their resting phenotype in blood. Circulating CD8$^+$ T cells were also largely characterized the upregulation of transcripts involved in the regulation of gene expression and translation processes (i.e. *EEF2, EIF3ESRP9, UBA52*), protein folding (*CCT3, VBP1, PFDN5*).

Upregulated genes in plaque CD8$^+$ T cells were involved in T cell activation (i.e. *HLA-A, HLA-DR*), chemotaxis (i.e. *CCL5, CCL4, CXCR6, XCL1*) and T cell exhaustion (i*.e. EOMES, PDCD1*) (**Supplementary Fig. 5e**). Signaling pathway analysis (**Supplementary Fig. 5f**) confirmed the co-existence of activated, pro-inflammatory and exhausted CD8$^+$ T cells in plaques. Top up-regulated pathways included IFNγ , AP-1, cytokine signaling and TRAF6 mediated induction of cytokines, chemotaxis and immune interactions, as well as PD-1, whose inhibitory signals play a major role during early stage of T cell activation as well as in overt T cell exhaustion during chronic inflammation and cancer[5,7,8].

Hierarchical clustering of CD8$^+$ T cells across plaque and blood identified 8 clusters with the majority (clusters 1, 2 and 4-7) being enriched in plaque and only two (clusters 3 and 8) in blood (**Supplementary Fig. 5g**). Signaling pathway analysis (**Supplementary Fig. 5h**) revealed heterogeneous and specialized states for each cluster. Clusters 1 and 7 were the more activated, with cluster 1 being more cytotoxic and expressing genes associated with T cell migration, and differentiation. Consistently, we found the activation of Rho, PCKθ  and NFAT signaling, all implicated in T cell activation[9], in cluster 7, and the exclusive upregulation of PCKθ signaling, which is required for TCR-induced T cell activation[10] in cluster 1. CD8$^+$ T cells of the other clusters were less active and cells of cluster 2 were uniquely characterized by the upregulation of the T cell exhaustion pathway. Overall these results confirm our CyTOF observations that like CD4$^+$ T cells, CD8$^+$ T cells in plaque present an activated, inflammatory phenotype.

***Single-Cell Proteomic Analysis of ASYM and SYM plaques***

We next analyzed the plaque-specific adaptive immune dysregulations from our T cell MetaClustering analysis. We found that the T cell MCs were not associated with either clinical characteristics or the type V and VI AHA plaque-type[11] seen in the prospectively enrolled patients of cohort 2 (**Supplementary Fig. 2d, f, h**).

Our analysis of the T cell MC frequencies determined that only MC12 was impacted by clinical phenotype. However, other plaque-enriched MCs displayed several differences in surface marker expression, suggesting plaque specific functional alterations associated with recent ischemic cerebrovascular events (**Supplementary Fig. 6c)**. In MC 10, SYM plaques displayed reduced CD103 expression, which is involved in maintaining tissue residency of TRM in tissues[12]. In MC 11, SYM plaques showed a trend towards reduced CD27, CD69 and GZMB expression. Although these changes did not reach statistical significance after correcting for multiple comparisons (CD27 and GZMB q value=0.06; CD69 q value=0.09), these data suggest a lower degree of activation, a more differentiated phenotype, and reduced cytolytic functions in plaques of SYM patients[13-16]. In the blood of SYM patients, MC 11 had significantly increased CD28 and CXCR3 levels and reduced CCR4 and GZMB levels, suggesting that these circulating cells are more activated than those of ASYM patients and are prone to migrating to the atherosclerotic vascular site. Finally, in MC 20, SYM plaques had a trend in increased levels of costimulatory marker CD28 (q value= 0.06), but no significant changes were identified for this MC in blood. These results suggest distinct functional variations in $CD8^+$ T cells (MC 10 and MC 11). Particularly, $CD8^+$ T cells of MC 11 displayed a less activated and more exhausted phenotype in SYM plaques, while mirroring cells of MC 11 in blood of the same SYM patients were activated and prone to infiltrate the atherosclerotic vascular tissue.

### SUPPLEMENTARY FIGURE LEGENDS

**Supplementary Figure 1. Interplay Between Cellular Populations.** (a) Similarity matrix of MC frequencies from n=15 patients. *indicates statistically significant correlations, p value<0.05 (b) Associations between the MC frequencies using the Spearman correlation method.

**Supplementary Figure 2. Pathological Characteristics of Atherosclerotic Plaques.** Pie charts of the atherosclerotic plaques used in the entire study (a) and segregated by cohorts (c, d) classified according to the American Heart Association criteria[84]. (b) Representative images of type V and type VI atherosclerotic lesions stained with Masson's trichrome stain. (d, g) Clustering of patients (columns) by MetaCluster frequencies (rows) from (d) Cohort 1 (discovery cohort, n=15), or (g) Cohort 2 (validation cohort, n=23). Top bars indicate the clinical characteristics (Sex,

Hypertensive, Diabetic, or BMI) and the associated AHA plaque type. Grey dendrogram bars below are identified clusters of patients. Clustering was performed using the average cosine distance using *Clustergrammer*. Individual clusters of patients were analyzed for their association with clinical characteristics and statistical significance (p values) is presented in the tables (e, h). P values were determined using the Binomial proportion test for each category in *Clustergrammer*.

**Supplementary Figure 3 Manual Gating Analysis of CYTOF data.** Manual gating analysis of the T-cell compartment from n=24 patients. (a) Representative viSNE plots of the distribution (from left to right) of blood and plaque, of CD4 and CD8 populations, and the distribution of T-cell subsets. (b) Population frequencies of cells gated for their effector status (naïve, CM, EM, and EMRA), or (c) classical definition (Th1, Th2, Th17, and Treg). For scatter bar plots, data were analyzed using the Wilcoxon test, *p<0.05, ***p<0.001, ****p<0.0001. Values are mean ± SD.

**Supplementary Figure 4. Tissue–specific T cell Metaclustering Identifies a Replicative Senescent $CD8^+$ T Cell population.** (a) Heatmap of MetaCluster communities of $CD3^+$ T Cells from atherosclerotic plaque by surface marker expression tissue (n=23). (b) Correlations between TCR clonality in plaque and the frequency of MetaCluster 25. (c) Representative viSNE plots showing the MC distribution and expression of selected markers. (d-e) Manually gated $CD8^+$ T cells from n=37 plaques (d) viSNE plot overlaid by $CD8^+CD127^+$ and $CD8^+$ $CD127^–$ populations. (e) Quantitation of subpopulation frequencies, assessed by paired Student's t test. (f) viSNE plots overlaid with expression of T cell functional markers. (g) Scatter plots of the median raw values of T cell functional markers. p values were determined using Wilcoxon test, *p<0.05, **p<0.01, ***p<0.001, ****p<0.0001. Values are mean ± SD. (h) Correlations between TCR clonality and $CD8^+CD127^-$ cell frequency. The Spearman statistic was used for the correlations for (b) and (h). (i) ViSNE plots of representative $CD8^+$ T cell population in plaque overlaid with CD127 expression (top), or by subpopulation: $CD127^+$ (blue), $CD127^–$ (yellow), and $Ki67^+$ (red) (bottom). (j) Percent of $Ki67^+$ cells in CD127 subpopulations. Statistic determined using the t test. (k) Representative contour plots of $CD8^+CD127^-$ plaque T Cells gated for CD57 expression, and the $CD8^+CD127^-CD57^+$ subpopulation gated for Ki67 expression.

**Supplementary Figure 5. Single-Cell Gene Expression analysis of $CD8^+$ and $CD4^+$ T Cells in paired blood and plaque.** Single-cell gene expression analysis of (a-d) $CD4^+$ and (e-h) $CD8^+$ T cells. (a, e) Volcano plot of the top 5000 Differentially Expressed Genes in plaque (purple) or blood (pink). (b, f) Pathway analysis of T cell gene upregulated in blood or plaque. Bars indicate the combined score from *Enrichr* (see methods). p values were determined using the Fisher's exact test, **p<0.01, ***p<0.0001, ****p<0.0001. (c, g) Top 50 variable genes in T cells across tissues hierarchically clustered. Rows indicate z-scored gene expression values, columns indicate individual cells. Categories above the heatmap indicate the cluster number, and below indicate the p values associated with the tissue enrichment. p values were determined using the Welch's t test with unequal variance, *p<0.05, **p<0.01, ***p<0.0001, ****p<0.0001. Adjacent boxes list key genes for cluster enrichment (g, i) Ingenuity Pathway Analysis (IPA) of the top 5000 DEGs per cluster in $CD4^+$ T cells (d) or $CD8^+$ T cells (h).

**Supplementary Figure 6. T cell Dysregulations between SYM and ASYM Immune Cells.** Bar charts of MC frequencies (a) and MC marker expression (b-c) in plaque enriched $CD4^+$ (b) or $CD8^+$ (c) T cell MCs. Blood (right) and plaque (left) of asymptomatic patients (A-SYM, blue bars) and symptomatic patients (SYM, red bars). Statistics were determined by multiple t-test and FDR (1%) correction using the two-stage step up procedure of Benjamini, Krieger, and Yekuteli.

**Supplementary Figure 7. Receptor-Ligand Paracrine Interactions Differentially Regulated between SYM and ASYM.** Differential regulation of top 30 differentially regulated in ASYM and SYM ligands paracrine ligand-receptor interactions between symptomatic and asymptomatic cells based on the absolute value sum of all ligand receptor interactions (see methods). (a) $CD4^+$ T cell driven interactions, (b) $CD8^+$ T cell interactions, and (c) macrophage interactions. Plots show the ligand as rows and receptor as columns. Ligands and receptors in the matrix are ordered based on sum in *Clustergrammer*.

**Supplementary Figure 8. Receptor-Ligand Autocrine Interactions Differentially Regulated between SYM and ASYM.** Differential regulation of top 30 differentially regulated in ASYM and SYM ligands autocrine ligand-receptor interactions between symptomatic and asymptomatic cells based on the absolute value sum of all ligand receptor interactions (see methods).  (a) $CD4^+$ T cell driven interactions, (b) $CD8^+$ T cell interactions, and (c) macrophage interactions. Plots show the ligand as rows and receptor as columns. Ligands and receptors in the matrix are ordered based on sum in *Clustergrammer*.

**Supplementary Figure 9. CITE-seq ADT and GEX relationships.** Correlations of Surface marker expression from ADT data and corresponding gene expression from the CITE-seq data.

**Supplementary Figure 10. Single-Cell RNA Sequencing Batch Correction.** UMAP dimensionality reduction plots are shown for T cells and Macrophages before and after batch correction. Cells from specific subjects are color coded.

### SUPPLEMENTARY TABLES

**Supplementary Table 1.** Patient Demographics and Clinical Characteristics

**Supplementary Table 2.** Fold Change and p Values of CYTOF MetaCluster Analyses

**Supplementary Table 3.** Antibodies used in this Study

**Supplementary Table 4.** Gating Strategy

**Supplementary Table 5.** Top 500 Differentially Expressed Genes in Blood versus Plaque Cells from CITE-seq Data

**Supplementary Table 6.** Top 500 Differentially Expressed Genes in SYM versus ASYM from scRNA-seq Data
